# Characterization of the exopolysaccharide biosynthesis pathway in *Myxococcus xanthus*

**DOI:** 10.1101/2020.06.08.139634

**Authors:** María Pérez-Burgos, Inmaculada García-Romero, Jana Jung, Eugenia Schander, Miguel A. Valvano, Lotte Søgaard-Andersen

## Abstract

*Myxococcus xanthus* arranges into two morphologically distinct biofilms depending on its nutritional status, i.e. coordinately spreading colonies in the presence of nutrients and spore-filled fruiting bodies in the absence of nutrients. A secreted polysaccharide referred to as exopolysaccharide (EPS) is a structural component of both biofilms and is also important for type IV pili-dependent motility and fruiting body formation. Here, we characterize the biosynthetic machinery responsible for EPS biosynthesis using bioinformatics, genetics, heterologous expression, and biochemical experiments. We show that this machinery constitutes a Wzx/Wzy-dependent pathway dedicated to EPS biosynthesis. Our data support that EpsZ (MXAN_7415) is the polyisoprenyl-phosphate hexose-1-phosphate transferase responsible for initiation of the repeat unit synthesis. Heterologous expression experiments support that EpsZ has galactose-1-P transferase activity. Moreover, MXAN_7416, renamed Wzx_EPS_, and MXAN_7442, renamed Wzy_EPS_, are the Wzx flippase and Wzy polymerase responsible for translocation and polymerization of the EPS repeat unit, respectively. Also, in this pathway, EpsV (MXAN_7421) is the polysaccharide co-polymerase and EpsY (MXAN_7417) the outer membrane polysaccharide export (OPX) protein. Mutants with single in-frame deletions in the five corresponding genes had defects in type IV pili-dependent motility and a conditional defect in fruiting body formation. Furthermore, all five mutants were deficient in type IV pili formation and genetic analyses suggest that EPS and/or the EPS biosynthetic machinery stimulates type IV pili extension. Additionally, we identify a polysaccharide biosynthesis gene cluster, which together with an orphan gene encoding an OPX protein make up a complete Wzx/Wzy-dependent pathway for synthesis of an unknown polysaccharide.

**Importance:** The secreted polysaccharide referred to as exopolysaccharide (EPS) has important functions in the social life cycle of *M. xanthus*; however, little is known about how EPS is synthesized. Here, we characterized the EPS biosynthetic machinery and show that it makes up a Wzx/Wzy-dependent pathway for polysaccharide biosynthesis. Mutants lacking a component of this pathway had reduced type IV pili-dependent motility and a conditional defect in development. Also, these analysis suggest that EPS and/or the EPS biosynthetic machinery is important for type IV pili formation.

## Introduction

Bacteria often exist in biofilms, which are surface-associated communities where cells are embedded in a self-produced extracellular matrix (1). Typically, this matrix is composed of proteins, extracellular DNA (eDNA) and polysaccharides (2). The polysaccharides serve several functions in a biofilm including structural roles, hydration, adhesion to substrates, cohesion between cells and protection against antibacterials, grazing and bacteriophages (2-4).

The Gram-negative delta-proteobacterium *Myxococcus xanthus* is a model organism to study social behaviors in bacteria. Depending on their nutritional status, *M. xanthus* cells organize into two morphologically distinct biofilms (5, 6). In the presence of nutrients, cells grow, divide and move across surfaces by means of two motility systems to generate colonies that are embedded in a polysaccharide referred to as exopolysaccharide (EPS) and in which cells at the colony edge spread outwards in a highly coordinated fashion (6-8). Under nutrient limitations, growth ceases and cells alter their motility behavior and begin to aggregate. The aggregation process culminates in the formation of mounds of cells inside which the rod-shaped cells differentiate into environmentally resistant spores leading to the formation of mature fruiting bodies (5, 6). EPS also makes up a substantial part of individual fruiting bodies (9-11).

The two motility systems of *M. xanthus* are important for formation of both biofilms (12). One motility system depends on type IV pili (T4P), which are highly dynamic filaments that undergo cycles of extension, surface adhesion and retraction. Retractions generate a force sufficient to pull a cell forward (13). The second system is for gliding motility and depends on the Agl/Glt complexes (6, 7). Generally, T4P-dependent motility involves movement of groups of cells while gliding motility involves the movement of individual cells (12, 14).

Besides its role as a structural component of spreading colonies and fruiting bodies, EPS in *M. xanthus* is also important for T4P-dependent motility (9, 15) and fruiting body formation (9, 10, 16-18). It has been proposed that EPS stimulates T4P-dependent motility by stimulating retraction of T4P (15, 19). Most insights into the function of EPS in *M. xanthus* have been obtained from analyses of regulatory mutants with altered levels of EPS synthesis. Among these mutants, the best studied include those of the Dif chemosensory system and the SgmT/DigR two component system. The Dif system is a key regulator of EPS synthesis; analyses of *dif* (previously *dsp* (10, 20, 21)) mutants have shown that decreased EPS accumulation (18, 21, 22) causes defects in T4P-dependent motility and fruiting body formation (17, 18). While the phosphotransfer reactions within the Dif system have been described in detail (22, 23), it is unknown how the Dif system stimulates EPS synthesis. Similarly, mutants of the SgmT/DigR system in which DigR is a DNA-binding response regulator have increased EPS accumulation, reduced T4P-dependent motility and a defect in fruiting body formation (24, 25). Transcriptome analyses support that this inhibitory effect is not caused by a direct effect on the expression of genes for EPS synthesis (25). Compared to the several identified regulators of EPS synthesis, relatively little is known about EPS biosynthesis. Here, we focused on the identification of proteins directly involved in EPS biosynthesis.

Synthesis of bacterial cell surface polysaccharides can occur via three different pathways, the Wzx/Wzy, the ABC-transporter or the synthase-dependent pathway (26, 27) (Fig. 1A). In the Wzx/Wzy and ABC-transporter dependent pathways, synthesis generally starts with the transfer of a sugar-1-P from a UDP-sugar to an undecaprenyl phosphate (Und-P) molecule in the inner leaflet of the inner membrane (IM) to form an Und-PP-sugar molecule (28). The priming enzymes are broadly classified in two groups, polyisoprenyl-phosphate hexose-1-phosphate transferases (PHPTs) or polyisoprenyl-phosphate *N*-acetylhexosamine-1-phosphate transferases (PNPTs) (29). Subsequently, the polysaccharide chain is elongated by the action of specific glycosyltransferases (GTs) and this depends on the specific pathway. In the Wzx/Wzy-dependent pathway, GTs synthesize the repeat unit of the polysaccharide on the cytoplasmic side of the IM; each unit is then translocated across the IM by the Wzx flippase and polymerized by the Wzy polymerase into a longer chain. Chain length is generally controlled by a Wzz protein, which belongs to the polysaccharide co-polymerase (PCP) family and results in the formation of polysaccharide molecules with a range of lengths (30, 31). By contrast, in the ABC-transporter dependent pathway, the full-length polysaccharide chain is synthetized on the cytoplasmic side of the IM, and is then translocated across the IM by an ABC transporter (32). In the synthase-dependent pathway, synthesis and translocation across the IM take place simultaneously by a multifunctional synthase protein complex in the IM (33). In the Wzx/Wzy and ABC-transporter dependent pathways, the polysaccharide molecule reaches the cell surface by translocation through an outer membrane (OM) polysaccharide export (OPX) protein and in the synthase-dependent pathway translocation occurs via an OM β-barrel protein (26, 33).

**Figure 1.**
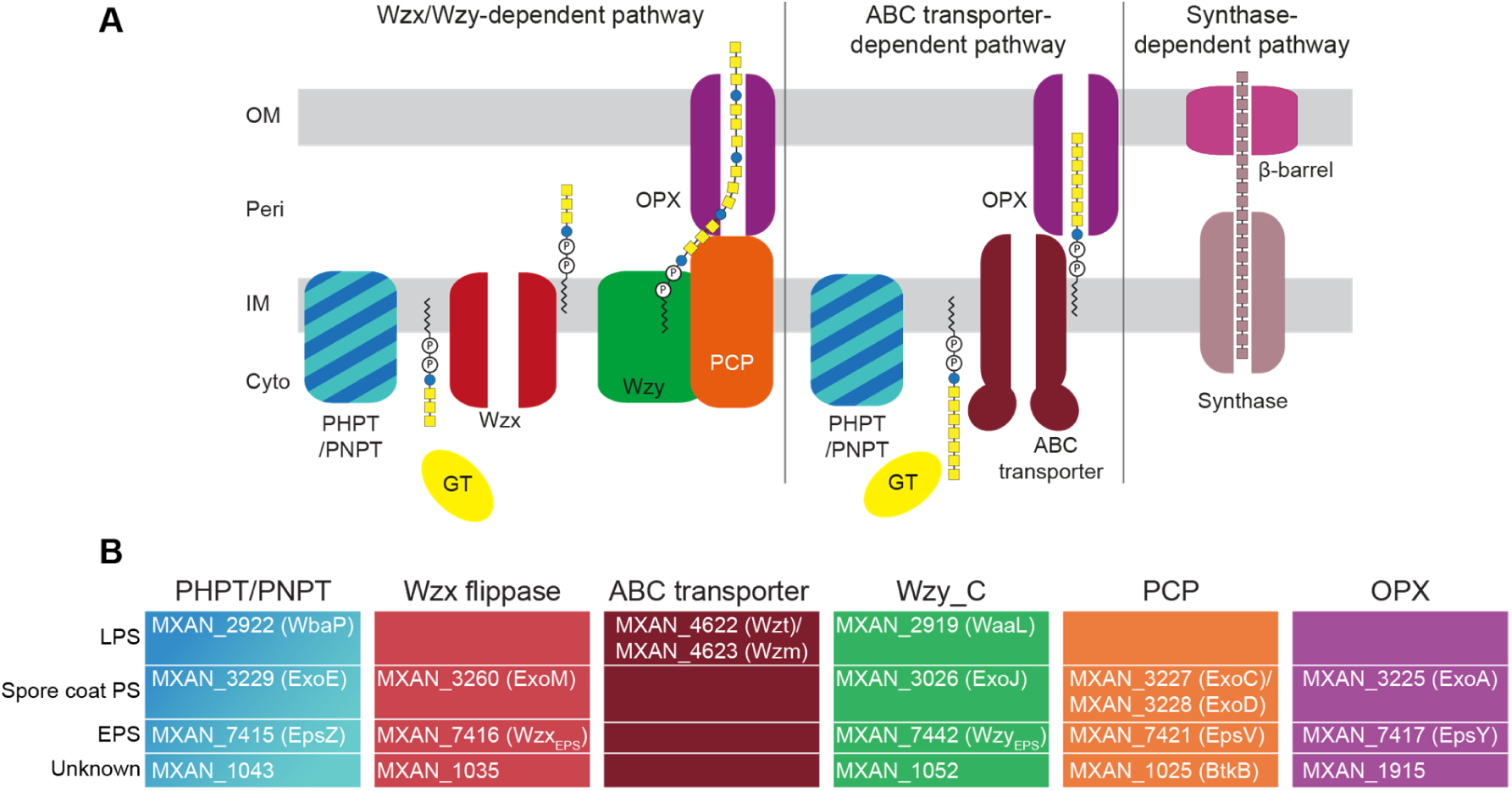
Identification of homologs of polysaccharide biosynthesis proteins in *M. xanthus.* (A) Schematic of the three pathways for polysaccharide biosynthesis in Gram-negative bacteria. (B) Bioinformatics-based identification of homologs of polysaccharide biosynthesis proteins in *M. xanthus.* Color code as in (A). Note that WaaL is the LPS O-antigen ligase (37) while the remaining three proteins with a Wzy_C domain are predicted polymerases.

The *eps* locus in *M. xanthus* was identified by transposon mutagenesis and shown to encode homologs of proteins involved in polysaccharide biosynthesis (9). Moreover, several *eps* genes were identified as essential for EPS biosynthesis (9, 22, 34-36). Here, we searched the re-annotated *eps* locus and the remaining *M. xanthus* genome for homologs of proteins for polysaccharide biosynthesis. We report that the *eps* locus encodes a complete Wzx/Wzy-dependent pathway for EPS biosynthesis. In-frame deletions in the corresponding genes specifically resulted in EPS biosynthesis defects without interfering with LPS O-antigen, spore coat polysaccharide and peptidoglycan (PG) biosynthesis. Phenotypic analysis of these mutants including complementation experiments demonstrated that they have a defect in T4P-dependent motility and conditional defects in development. In addition, we identify a polysaccharide biosynthesis gene cluster of unknown function that together with an orphan gene encoding an OPX protein encode a complete Wzx/Wzy-dependent pathway for biosynthesis of a polysaccharide of unknown function.

## Results

### Identification of homologs of proteins of Wzx/Wzy-dependent pathways for polysaccharide biosynthesis and export

To identify genes for EPS biosynthesis, we searched the *M. xanthus* genome for homologs (*see Materials and Methods*) of the membrane components of the three biosynthesis pathways (Fig. 1A). We identified homologs encoding predicted proteins of the Wzx/Wzy and ABC-transporter pathways but none corresponding to a synthase-dependent pathway (Fig. 1B). Several of these homologs were previously shown to be important for LPS synthesis or spore coat polysaccharide biosynthesis (37-41) (Fig. 1B). Notably, none of these proteins are required for EPS biosynthesis. The MraY homolog (MXAN_5607), which belongs to the PNPT family and is involved in PG synthesis, was not considered here.

The reannotated *eps* locus consists of two gene clusters (*MXAN_7515-_7422* and *MXAN_7441-_7451*) that encode all the proteins of a complete Wzx/Wzy-dependent pathway (Fig. 2A, Table S1). Specifically, these two gene clusters encode homologs of a PHPT (EpsZ/MXAN_7415), a Wzx flippase (MXAN_7416), a Wzy polymerase (MXAN_7442, previously SgnF (42)), a PCP protein (EpsV/MXAN_7421) and an OPX protein (EpsY/MXAN_7417) as well as five GTs (EpsU/MXAN_7422, EpsH/MXAN_7441, EpsE/MXAN_7445, EpsD/MXAN_7448, EpsA/MXAN_7451) and a serine O-acetyltransferase (EpsC/MXAN_7449). Previous genetic analyses using transposon insertions, plasmid insertions or in-frame deletion mutants demonstrated that genes in both clusters are important for EPS synthesis (9, 22, 34, 35) (Fig. 2A). Also, genes in both clusters were previously shown to be important for T4P-dependent motility without directly testing for EPS synthesis (42) (Fig. 2A). The two gene clusters are separated by 13 genes encoding proteins predicted not to be directly involved in polysaccharide synthesis (Fig. 2A and Table S1). Consistently, genetic analyses for some of these genes confirmed that they are not important for EPS synthesis (9) except for *MXAN_7440* (Nla24/EpsI), which encodes a c-di-GMP binding NtrC-like transcriptional regulator (36, 43) that is phosphorylated by the histidine kinase MXAN_7439 (44).

**Figure 2.**
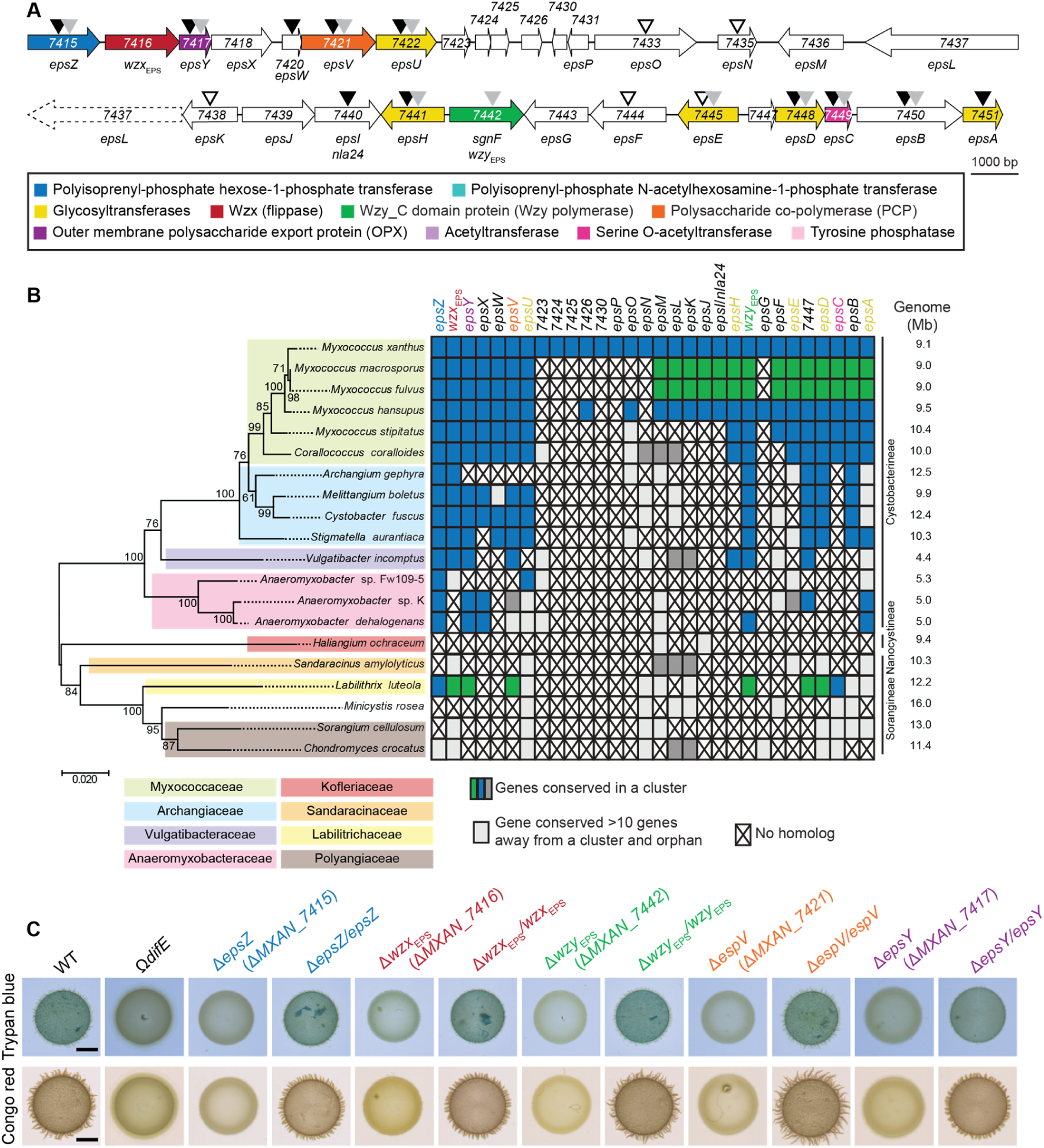
Bioinformatics and genetic analysis of the *eps* locus. (A) *eps* locus in *M. xanthus*. Genes are drawn to scale and MXAN number or gene name is indicated (Table S1). The color code indicates predicted functions as indicated in the key and are used throughout. Black, grey and white arrow heads indicate mutations previously reported to cause a defect in EPS synthesis (black) (9, 22, 34, 36), a defect in T4P-dependent motility but with no test of EPS synthesis (42)(grey) and no effect on EPS synthesis (9). (B) Taxonomic distribution and synteny of *eps* genes in Myxococcales with fully sequenced genomes. A reciprocal best BlastP hit method was used to identify orthologs. 16S rRNA tree of Myxococcales with fully sequenced genomes (left). Genome size, family and suborder classification are indicated (right). To evaluate gene proximity and cluster conservation, 10 genes were considered as the maximum distance for a gene to be in a cluster. Genes found in the same cluster (within a distance of <10 genes) are marked with the same color (i.e. blue, green and dark grey). Light grey indicates a conserved gene that is found somewhere else on the genome (>10 genes away from a cluster); a cross indicates no homolog found. (C) Determination of EPS synthesis. 20 μl aliquots of cell suspensions of strains of the indicated genotypes at 7 × 10^9^ cells ml^-1^ were spotted on 0.5% agar supplemented with 0.5% CTT and Congo red or Trypan blue and incubated 24 h. In the complementation strains, the complementing gene was expressed ectopically from the native (*epsZ, wzx*_EPS_ and *wzy*_EPS_) or the *pilA* promoter on a plasmid integrated in a single copy at the Mx8 *attB* site. The Ω*difE* mutant served as a negative control.

In a bioinformatics approach searching for orthologs of the proteins encoded by the entire *eps* locus in all fully sequenced Myxococcales genomes using a reciprocal best BlastP hit method as in (41), we found that the two gene clusters encoding proteins for polysaccharide synthesis are largely conserved in closely related Cystobacterineae (Fig. 2B). Importantly, in several of these genomes, the two clusters are present in a single uninterrupted gene cluster (Fig. 2B). Interestingly, in *M. macrosporus* and *M. fulvus*, the two gene clusters are separated by a set of genes that are conserved between these two organisms but not homologous to the genes separating the two clusters in *M. xanthus*. Together, based on previous gene genetic analyses and because genes for polysaccharide biosynthesis are often clustered (45), our data support that the two separated gene clusters in the *M. xanthus eps* locus encode for a Wzx/Wzy-dependent pathway for EPS synthesis.

We also identified a second locus encoding homologs of a Wzx/Wzy pathway (Fig. 3A; Table S2). Specifically, this locus encodes homologs of a PNPT (MXAN_1043), a Wzx flippase (MXAN_1035), a Wzy polymerase (MXAN_1052), a Wzc chain length regulator (MXAN_1025 or BtkB (46)) of the PCP-2 family as well as 10 GTs (MXAN_1026, MXAN_1027, MXAN_1029, MXAN_1030, MXAN_1031, MXAN_1032, MXAN_1033, MXAN_1036, MXAN_1037, MXAN_1042) and two acetyltransferases (MXAN_1041 and MXAN_1049). Finally, we identified a gene encoding an OPX protein (MXAN_1915) that is not part of a gene cluster encoding proteins involved in polysaccharide synthesis (Fig. 3B, Table S2). Using bioinformatics, as described above, we found that the large gene cluster as well as *MXAN_1915* are conserved in closely related Cystobacterineae (Fig. 3C). Importantly, the *MXAN_1915* ortholog of *Sandaracinus amylolyticus* is encoded in a cluster with homologs of *MXAN_1025, MXAN_1043* and *MXAN_1052*. Because the *MXAN_1025-_1052* locus does not encode an OPX homolog, these observation support that MXAN_1915 could function together with the proteins encoded by this locus and together they would make up a complete Wzx/Wzy-dependent pathway for biosynthesis of a polysaccharide. Based on these analyses, we hypothesized that the proteins encoded by the *eps* locus and the proteins encoded by the *MXAN_1025-_1052*/*_1915* loci make up two independent and dedicated pathways for polysaccharide synthesis.

**Figure 3.**
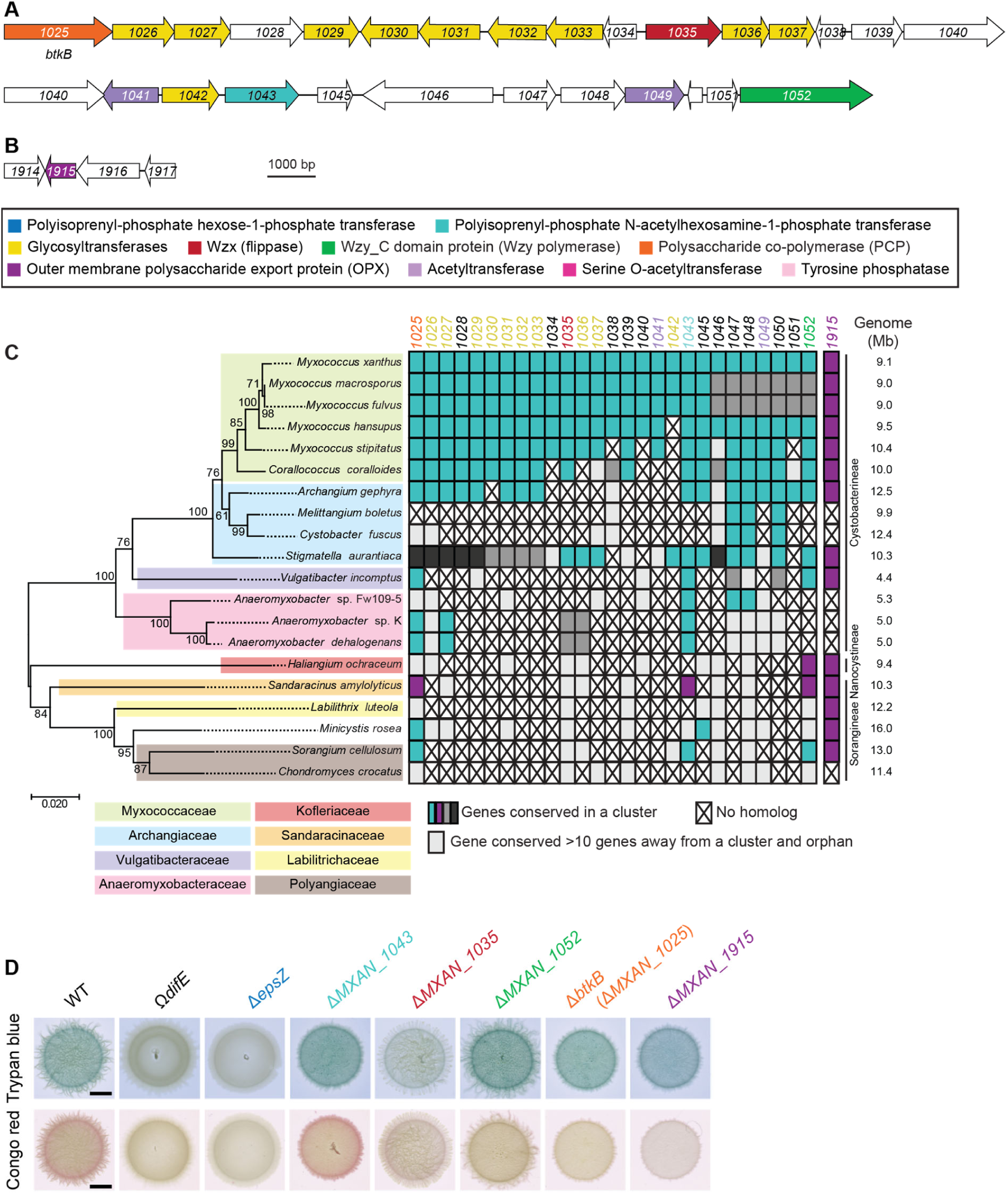
Bioinformatics and genetic analysis of the *MXAN_1025-1052/_1915* loci. (A, B) *MXAN_1025-1052 and _1915* loci in *M. xanthus*. Genes are drawn to scale and MXAN number or gene name is indicated (Table S2). The color code indicates predicted functions as indicated in the key and are used throughout. (C) Taxonomic distribution and synteny of genes in the *MXAN_1025-1052/_1915* loci in Myxococcales with fully sequenced genomes. A reciprocal best BlastP hit method was used to identify orthologs. 16S rRNA tree of Myxococcales with fully sequenced genomes (left). Genome size, family and suborder classification are indicated (right). To evaluate gene proximity and cluster conservation, 10 genes were considered as the maximum distance for a gene to be in a cluster. Genes found in the same cluster (within a distance of <10 genes) are marked with the same color (i.e. cyan, magenta, and dark and middle dark grey). Light grey indicates a conserved gene that is found somewhere else on the genome (>10 genes away from a cluster); a cross indicates no homolog found. (C) Determination of EPS synthesis. 20 μl aliquots of cell suspensions of strains of the indicated genotypes at 7 × 10^9^ cells ml^-1^ were spotted on 0.5% agar supplemented with 0.5% CTT and Congo red or Trypan blue and incubated 24 h. The Ω*difE* mutant served as a negative control. Scale bars, 3 µm.

### The *eps* locus is essential for EPS biosynthesis

To test for the importance of genes of the *eps* locus and the *MXAN_1025-_1052/_1915* loci for EPS synthesis, we generated ten in-frame deletions in genes encoding the five conserved core components of Wzx/Wzy-dependent pathways (i.e. the genes for the PH/NPT, Wzx, Wzy, PCP and OPX). Subsequently, we used plate-based colorimetric assays with either Congo red or Trypan blue to assess EPS biosynthesis. As a negative control, we used a Ω*difE* mutant, which has a defect in EPS synthesis (17).

All five mutations in the *eps* locus abolished EPS synthesis (Fig. 2C). Importantly, the EPS synthesis defects of these five Δ*eps* mutants were complemented by ectopic expression of the relevant full-length gene from a plasmid integrated in a single copy at the Mx8 *attB* site (Fig. 2C). By contrast, in case of the five in-frame deletions in the genes of the *MXAN_1025-_1052*/*_1915* loci, only the Δ*MXAN_1035* mutant, which lacks a Wzx flippase homolog (Fig. 3A-C) caused a significant decrease in EPS synthesis. Based on several arguments, we do not think that MXAN_1035 is directly involved in EPS synthesis but rather that the Δ*MXAN_1035* mutation results in titration of Und-P. First, mutation of *MXAN_7416*, which encodes a Wzx flippase homolog in the *eps* locus, completely blocked EPS synthesis (Fig. 2A-C) supporting that MXAN_7416 is the flippase involved in EPS biosynthesis. Second, as mentioned, enzymes of the same polysaccharide biosynthesis and export pathway are typically encoded in the same locus (45); however, the three other mutations in the *MXAN_1025-_1052* locus did not have a significant effect on EPS biosynthesis (Fig. 3D). Third, blocking translocation of a specific sugar unit across the IM can cause sequestration of Und-P and, thereby, result in pleiotropic effects on the synthesis of other polysaccharides (47-51). Consistently, a Δ*MXAN_1035* mutation was previously shown to cause a reduction in glycerol-induced sporulation (see below) likely by interfering with spore coat polysaccharide biosynthesis (39); however, MXAN_3260 (ExoM) was recently shown to be the flippase involved in spore coat polysaccharide synthesis (41). Together, these considerations support that it is unlikely that MXAN_1035 is part of the EPS biosynthesis machinery. In total, our results suggest that the *eps* locus encodes homologs of a Wzx/Wzy-dependent pathway for EPS biosynthesis. Therefore, we renamed MXAN_7416 and MXAN_7442 to Wzx_EPS_ and Wzy_EPS_. From here on, we focused on the five core components of the Wzx/Wzy-dependent pathway for EPS synthesis.

### Lack of EPS biosynthetic proteins does not affect spore coat polysaccharide, LPS synthesis or cell morphology

In addition to EPS, *M. xanthus* synthetizes O-antigen LPS (52) and a spore coat polysaccharide (53). As mentioned, because blocking synthesis of one polysaccharide can affect synthesis of other polysaccharide including PG by sequestration of Und-P through accumulation of Und-PP intermediates, we determined whether lack of the EPS biosynthetic proteins affects spore coat polysaccharide, LPS or PG synthesis.

Synthesis of the spore coat polysaccharide is essential for sporulation in *M. xanthus* (40, 54). To evaluate whether the Δ*eps* mutants synthetized spore coat polysaccharide, we analyzed sporulation independently of starvation. For this, we profited from an assay in which sporulation occurs rapidly and synchronously and is induced chemically by the addition of glycerol at a high concentration (0.5 M) to cells growing in nutrient-rich broth (55). In response to adding glycerol, cells of WT and all *eps* five in-frame deletion mutants rounded up during the first 4 h and had turned into phase-bright resistant spores by 24 h (Fig. 4A). Cells of the Δ*exoE* mutant, which lacks the PHPT for initiating spore coat polysaccharide biosynthesis and were used as a negative control (39, 41), remained rod-shaped and did not form phase-bright spores. Interestingly, the sporulation efficiency of all five *Δeps* mutants was increased compared to WT (Fig. 4A). Because the spores formed by WT under high concentrations of glycerol adhere to glass surfaces and each other forming large aggregates while the spores formed by the Δ*eps* mutants do not, we speculate that the ease of harvesting the EPS^-^ spores rather than the *eps* mutations *per se* results in an apparent increase in the overall sporulation efficiency. We conclude that lack of the EPS biosynthetic proteins does not cause a sporulation defect, in agreement with previous observations that mutation of *epsV* did not affect glycerol-induced sporulation (39).

**Figure 4.**
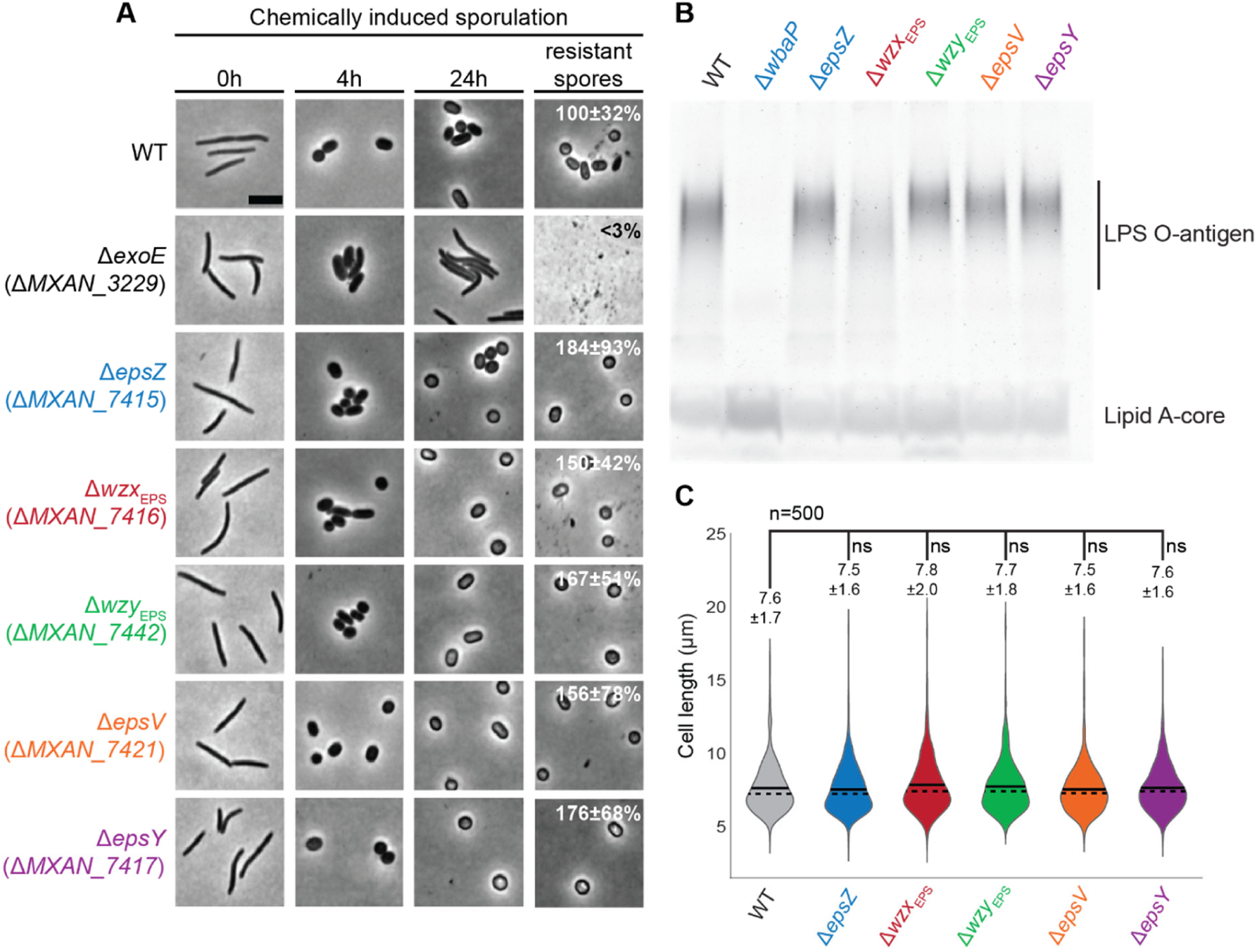
Phenotypic characterization of the Δ*eps* mutants. (A) Chemically induced sporulation. Sporulation was induced by addition of glycerol to a final concentration of 0.5 M. At 0, 4 and 24 h after glycerol addition, cell morphology was documented. In images labelled resistant spores, cells were exposed to sonic and heat treatment before microscopy. Sporulation frequency after sonic and heat treatment is indicated as the mean of three biological replicates relative to WT ± standard deviation. Scale bars, 5 µm. (B) Extracted LPS from the same number of cells was separated by SDS-PAGE and detected with Pro-Q Emerald 300. (C) Cell length measurements of the Δ*eps* mutants. Cell length is shown in a violin plot, which indicates the probability density of the data at different cell length values. *n* = 500 combined from two biological replicates and mean and median values are represented by a continuous and dashed line, respectively. Samples were compared using a Mann-Whitney test; ns, not significant.

LPS in total cell extracts was detected by Emerald staining and the Δ*wbaP* mutant, which lacks the PHPT for O-antigen biosynthesis, served as a negative control (37). A fast running lipid-A core band and polymeric LPS O-antigen bands were detected in LPS preparations of WT and the five Δ*eps* mutants, while only the lipid-A core band was detected in the Δ*wbaP* mutant (Fig. 4B). The Δ*wzx*_EPS_ mutant accumulated lower levels of LPS O-antigen (Fig. 4B). O-antigen in *M. xanthus* is synthesized via an ABC transporter-dependent pathway and lack of the Wzm/Wzt ABC transporter blocks LPS O-antigen synthesis (37, 38), suggesting that the Wzx_EPS_ is not directly involved in O-antigen synthesis. Therefore, we speculate that the reduced O-antigen level in the Δ*wzx*_EPS_ mutant could be caused by sequestration of Und-PP-linked EPS intermediates unable to be translocated across the membrane, which would reduce the available pool of Und-P for O-antigen synthesis.

Interference with PG synthesis during growth in *M. xanthus* causes morphological defects (56-58). Therefore, we used cell morphology as a proxy for PG synthesis to test whether lack of the EPS biosynthetic proteins interferes with PG synthesis during growth. Because, cell morphology and cell length of the five Δ*eps* mutants were similar to that of WT cells, we conclude that PG synthesis was not affected in the Δ*eps* mutants (Fig.4A (0 h) and 4C). Therefore, the proteins for EPS biosynthesis are not important for spore coat polysaccharide, LPS synthesis and PG synthesis, supporting that these proteins make up a pathway dedicated to EPS synthesis.

### MXAN_7415 has Gal-1-P transferase activity

EpsZ is the predicted PHPT of the EPS biosynthesis pathway. Similar to WcaJ_Ec_ from *E. coli* and WbaP_Se_ from *Salmonella enterica* (41, 59, 60), we identified a PF13727 (CoA_binding _3) domain, a C-terminal PF02397 (Bac_transf) domain and five transmembrane regions in EpsZ (Fig. 5A), all features of PHPT proteins. The fifth TMH of WcaJ_Ec_ does not fully span the IM and this results in the cytoplasmic localization of the C-terminal catalytic domain. This depends on residue P291 that causes a helix-break-helix in the structure and forms part of a DX_12_P motif conserved among PHPTs (60). Because EpsZ contains the DX_12_P motif and all the conserved essential residues important for catalytic activity that have been identified in the C-terminal catalytic region of WbaP (61) (Fig. 5B, Fig S1), we suggest that EpsZ is a PHPT with a membrane topology similar to that of WcaJ.

**Figure 5.**
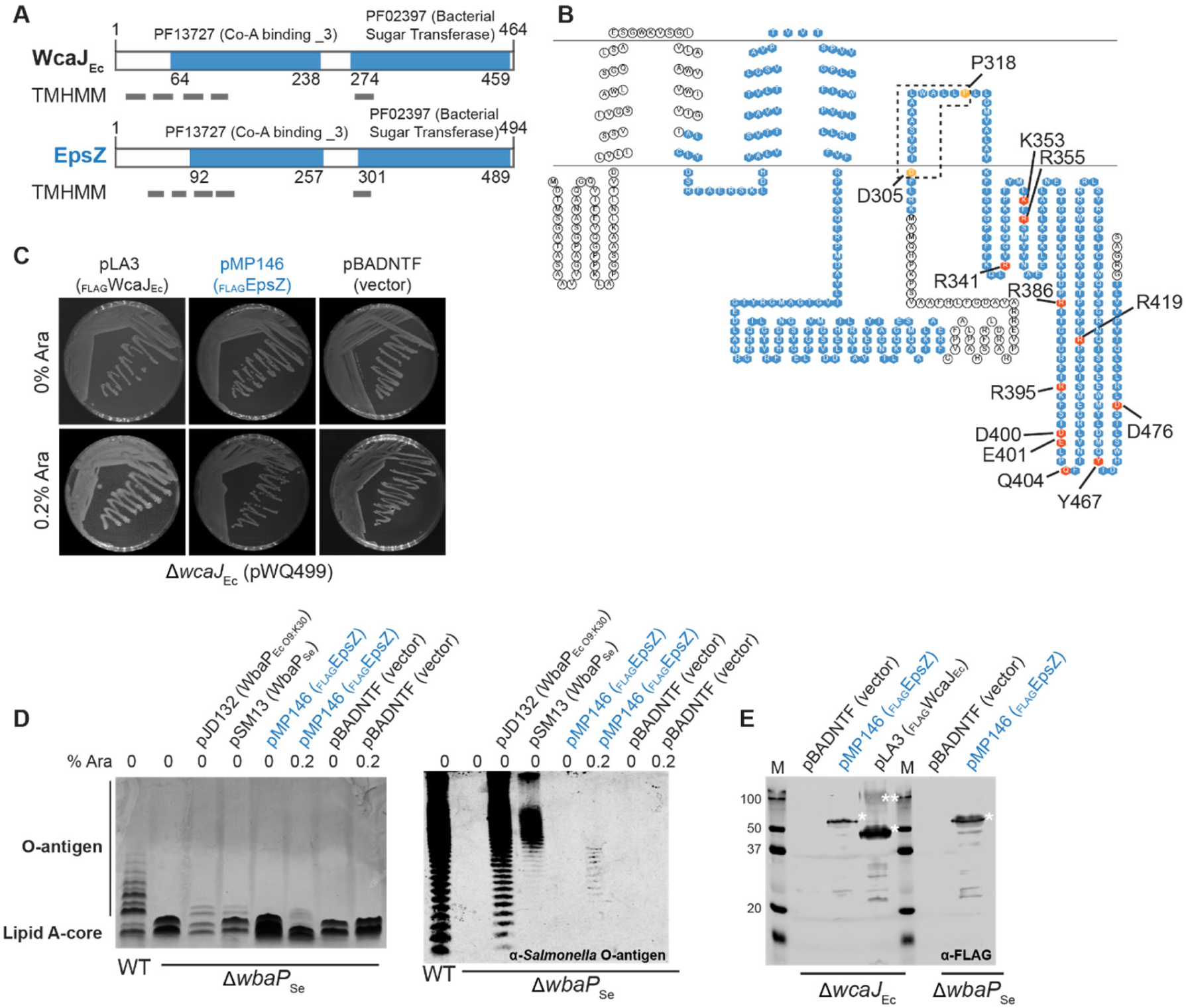
PHPT activity of MXAN_7415. (A) Domain and TMH prediction of EpsZ (MXAN_7415) and WcaJ of *E. coli* (WcaJ_Ec_). Grey rectangles indicate TMH. Numbers indicate domain borders. (B) Topology predictions for EpsZ (MXAN_7415). Domains are indicated in blue and conserved amino acids important for structure or activity of the protein are marked with orange and red, respectively. Sequence alignment of EpsZ (MXAN_7415) with WbaP_Se_, is shown in Fig. S1. (C-E) Complementation of colanic acid synthesis and LPS O-antigen in *E. coli* K-12 W3110 (Δ*wcaJ*_Ec_) and *S. enterica* LT2 (Δ*wbaP*_Se_) mutants, respectively by plasmids encoding the indicated PHPT proteins. (C) The *E. coli* Δ*wcaJ*_Ec_ mutant XBF1 containing pWQ499 (RcsA^+^) and the indicated complementing plasmids or vector control on LB plates was incubated overnight at 37°C with 10 μg ml^-1^ tetracycline (to maintain pWQ499) and with or without arabinose (Ara) to induce gene expression. Incubation was extended to 24-48 h at room temperature to further increase colanic polysaccharide synthesis. (D) Complementation of *S. enterica* Typhimurium LT2 Δ*wbaP*_Se_ mutant containing the indicated plasmids. LPS samples were extracted, separated by electrophoresis on SDS–14% polyacrylamide gels and silver stained (left panel) or examined by immunoblotting using rabbit *Salmonella* O antiserum group B (right panel). Each lane corresponds to LPS extracted from 10^8^ cells. Cultures included addition of arabinose as indicated. (E) Immunoblot using α-FLAG monoclonal antibody to confirm expression of _FLAG_MXAN_7415 and _FLAG_WcaJ in the Δ*wcaJ* mutant, and the expression of _FLAG_MXAN_7415 in *S. enterica*. Note that WbaP expressed from pSM13 was not tested since it is not fused to a FLAG tag. * and ** denote the monomeric and oligomeric forms of the PHPT proteins, usually present under the gel conditions required to ensure their visualization.

PHPTs generally utilize UDP-glucose (UDP-Glc) or UDP-galactose (UDP-Gal) as substrates to transfer Glc-1-P or Gal-1-P, respectively to Und-P (29, 62). Therefore, following the same strategy as previously reported (37, 41, 63), we tested whether EpsZ could functionally replace WcaJ_Ec_ or WbaP_Se_, which catalyse the transfer of Glc-1-P and Gal-1-P to Und-P, respectively. To this end, *epsZ* was cloned into pBADNTF resulting in plasmid pMP146, which encodes EpsZ with an N-terminal FLAG-tag (_FLAG_EpsZ) to facilitate detection by immunoblot and under the control of an arabinose inducible promoter.

WcaJ_Ec_ initiates colanic acid biosynthesis, which results in a strong glossy and mucoid phenotype of *wcaJ*_Ec_^+^ cells containing the plasmid pWQ499 encoding the positive regulator RcsA (60). An *E. coli* Δ*wcaJ*_Ec_ (pWQ499) mutant is complemented with the plasmid pLA3 in the presence of arabinose (60), which encodes _FLAG_WcaJ_Ec_ under the control of the arabinose inducible promoter (Fig. 5C). By contrast, no complementation was observed by _FLAG_EpsZ or the empty pBADNTF vector in the presence of arabinose (Fig. 5C), suggesting that EpsZ does not have Glc-1-P transferase activity.

WbaP_Se_ initiates O-antigen synthesis in *S. enterica* and the O-antigen synthesis defect of a Δ*wbaP*_Se_ mutant can be partially corrected by complementation with the plasmid pJD132, which encodes the *E. coli* O9:K30 WbaP_Se_ homolog (WbaP_Ec O9:K30_), and with the plasmid pSM13, which encodes WbaP_Se_ (59) (Fig. 5D, left panel). The differences in the O-antigen profile between the different complementation strains are likely due to different processing of the O-antigen as previously reported (59). Expression of _FLAG_EpsZ in the Δ*wbaP*_Se_ mutant in the presence of arabinose provoked a change of the LPS profile (Fig. 5D, left panel), while the empty pBADNTF vector did not affect the LPS profile. Because the effect of _FLAG_EpsZ on the O-antigen profile of the Δ*wbaP*_Se_ mutant was relatively modest by silver staining, we repeated these experiments using *Salmonella* O-antigen rabbit antibodies. As shown in Fig. 5D, right panel, also in this analysis, _FLAG_EpsZ complemented the Δ*wbaP*_Se_ mutant in the presence of arabinose. To test for accumulation of _FLAG_EpsZ in the *E. coli* and *S. enterica* strains when grown in the presence of arabinose, we performed immunoblots using α-FLAG antibodies (Fig. 5E). EpsZ accumulated in both strains predominantly in the monomeric form. By contrast, _FLAG_WcaJ_Ec_ showed the characteristic oligomeric and monomeric bands as previously reported for PHPTs (59). We conclude from these experiments that WbaP_Mx_ can transfer Gal-1-P onto Und-P.

### EPS or EPS biosynthetic proteins is important for T4P-dependent motility and T4P formation

Next, we tested the five Δ*eps* mutants for motility defects. To this end, cells were spotted on 0.5% and 1.5% agar, respectively (14). On 0.5% agar, WT cells formed the long flares characteristic of T4P-dependent motility while on 1.5% agar, WT displayed the single cells at the colony edge characteristic of gliding motility. The Δ*pilA* mutant, which lacks the major pilin subunit and does not assemble T4P (64), and the Δ*aglQ* mutant, which lacks a component of the gliding motility machinery (65, 66), were used as negative controls for T4P-dependent and gliding motility, respectively. As expected, the Δ*eps* mutants had a T4P-dependent motility defect forming colonies with shorter flares than WT as did the Δ*aglQ* mutant (Fig. 6A). The motility defects of the Δ*eps* mutants were complemented by ectopic expression of the relevant genes (Fig. 6A). On 1.5% agar, the Δ*eps* mutants displayed the single cell motility characteristic of gliding motility but the Δ*aglQ* mutant did not (Fig. 6A). Also, the total colony expansion was reduced similarly to the Δ*pilA* mutant. The reduced colony expansion was corrected in the five complementation strains (Fig. 6A). Because the Δ*aglQ* mutant made shorter flares on 0.5% agar and had no single cell motility on 1.5% agar, the Δ*pilA* mutant made no flares on 0.5% agar and had reduced colony expansion on 1.5% agar, while the five Δ*eps* mutants generated shorter flares on 0.5% agar and still had single cell motility on 1.5% agar, we conclude that lack of any single one of the five EPS biosynthetic proteins cause a defect in T4P-dependent motility but not in gliding motility.

**Figure 6.**
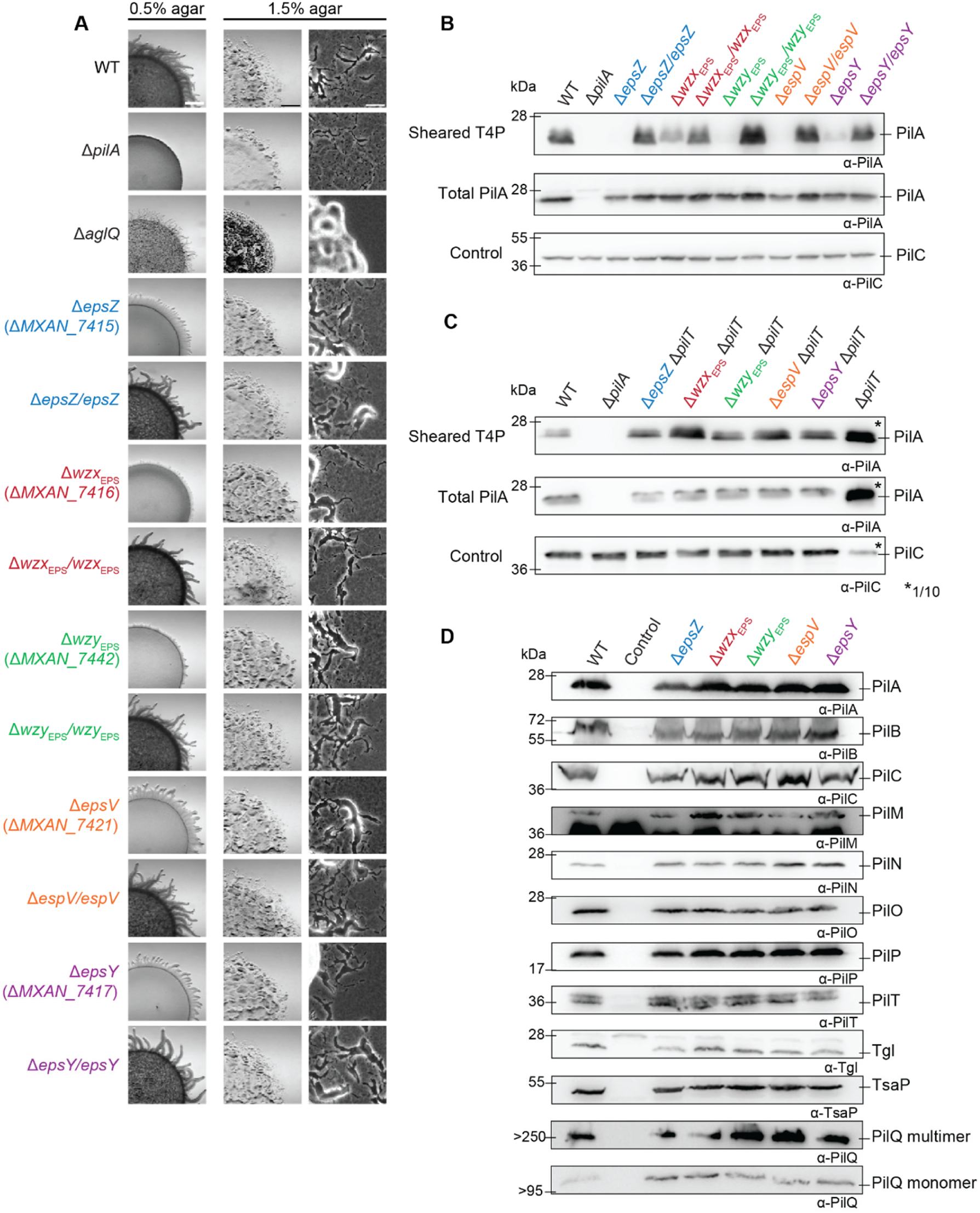
Motility analyses of Δ*eps* mutants. (A) Colony-based motility assay of Δ*eps* mutants. T4P-dependent motility and gliding motility were tested on 0.5% and 1.5% agar, respectively. Images were recorded after 24 h. Scale bars, 1 mm, 1 mm and 50 µm (left to right). (B-C) T4P shear off assay. Immunoblot detection of the major pilin PilA in sheared T4P (top) and in total cell extract (middle), where the same number of cells grown on 1% CTT 1.5% agar was loaded per lane. The top and middle blots were probed with α-PilA antibodies (calculated molecular mass 23.4 kDa). The middle blot was stripped and probed with antibodies against PilC (calculated molecular mass 45.2 kDa), as a loading control. (D) Immunoblot detection of proteins of the T4P machinery using α-PilA, -B, -C, -M, -N, -O, -P, -Q, -T, Tgl and TsaP antibodies. The same number of cells coming from exponentially growing liquid cultures was loaded per lane. As a negative control, cells containing a single in-frame deletion mutation in the relevant gene were used.

To understand the mechanism underlying the defect in T4P-dependent motility in the Δ*eps* mutants, we determined the level of T4P formation using a shear-off assay in which T4P are sheared off the cell surface and then the level of PilA assessed by immunoblotting. The PilA level in the sheared fraction was strongly reduced in all five Δ*eps* mutants while the total cellular level of PilA was generally similar to that in WT suggesting that these mutants have fewer T4P than WT cells (Fig. 6B). Because a reduced level of T4P can result from an extension defect or hyper-retraction, we deleted the *pilT* gene encoding the PilT retraction ATPase (67) in the five Δ*eps* mutants and then repeated the shearing assay. All five strains with the additional Δ*pilT* mutation assembled T4P at a higher level than the *pilT*^+^ strains, but at a significantly lower level than the Δ*pilT* strain (Fig. 6C). Thus, the five Δ*eps* mutants have a defect in T4P extension. Of note, the observation that the Δ*eps pilT*^+^ strains make fewer T4P than the Δ*eps* Δ*pilT* strains support that T4P still retract in the absence of the EPS biosynthetic machinery and/or EPS. These observations are in stark contrast to the observations for the Δ*difA* mutant, which lacks the MCP component of the Dif system and is strongly reduced in EPS synthesis (21). This mutant was reported to make T4P at WT levels (21) or to be hyper-piliated (15) and EPS was reported to stimulate T4P retractions in this mutant (15, 19). We conclude that lack of an EPS biosynthetic protein and/or EPS causes a reduction in T4P extension but the fewer T4P made can still retract.

To analyze whether the reduced T4P formation in the Δ*eps* mutants was caused by reduced synthesis of one or more of the 10 core proteins of the T4P machine (13, 68) or the Tgl pilotin (69), we determined their accumulation level in the five *eps* mutants. All 11 proteins were detected at WT levels in the Δ*eps* mutants (Fig. 6D). We conclude that the EPS biosynthetic machinery and/or EPS is important for T4P extension and, therefore, T4P-dependent motility.

Cell-cell cohesion has been suggested to depend on EPS (10, 16, 70). To evaluate whether the Δ*eps* mutants were affected in cell-cell cohesion and agglutination, we transferred exponentially growing cells to a cuvette and measured the change in cell density over time. WT cells agglutinated and sedimented during the course of the experiment, causing a decrease in the absorbance (Fig. 7). The Ω*difE* and mutant was used as a negative control and did not agglutinate over time (21). None of the five Δ*eps* in-frame deletion strains agglutinated (Fig. 7) and the agglutination defect was complemented in the complementation strains (Fig. 7).

**Figure 7.**
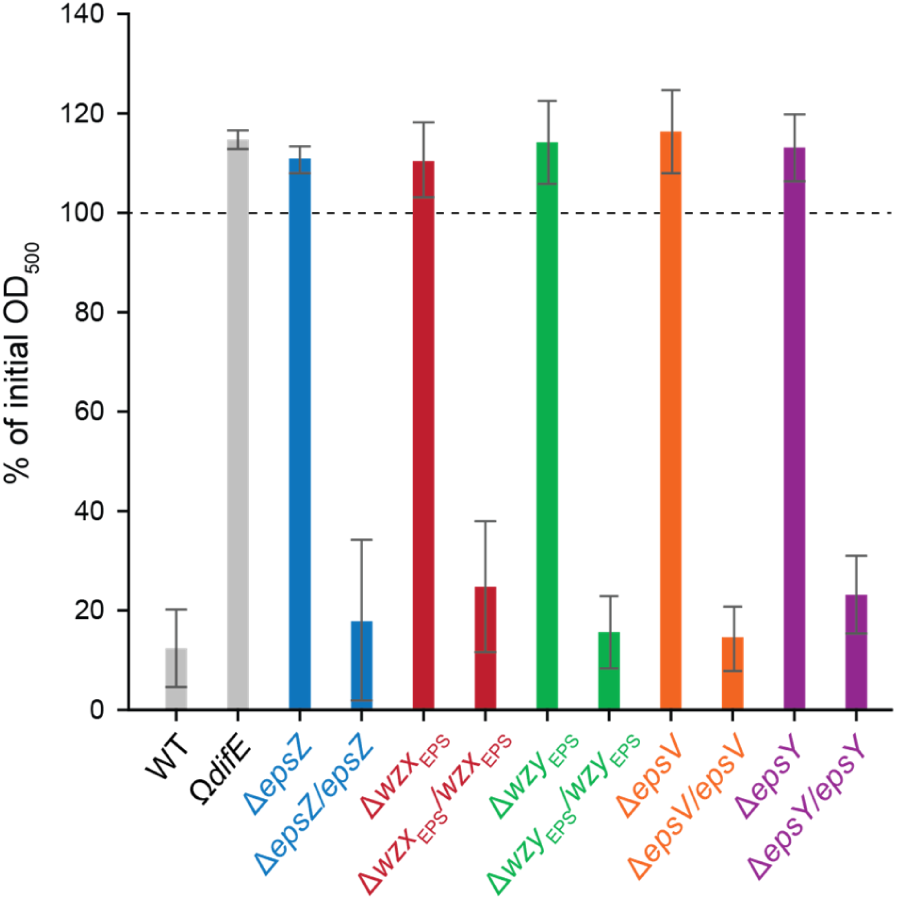
Analysis of Δ*eps* mutants for cell-cell cohesion and agglutination. Cell agglutination assay. 1 ml of exponentially growing cells were transferred to a cuvette. Agglutination was monitored by measuring the decrease in absorbance at 550 nm at 3 h relative to the initial absorbance for each strain. The graph shows data from three biological replicates as mean ± standard deviation.

### EPS or the Eps biosynthetic machinery is conditionally important for fruiting body formation

Next, we tested the five Δ*eps* mutants for development. On TPM agar and in submerged culture, WT cells had aggregated to form darkened mounds at 24 h of starvation. (Fig. 8). On TPM agar, the Δ*eps* mutants showed a delay in aggregation but eventually formed larger and less compact fruiting bodies and sporulated with an efficiency similar to that of WT (Fig. 8). Under submerged conditions, the Δ*eps* mutants did not aggregate to form fruiting bodies sporulation as expected from the cell-cell cohesion and agglutination defects and were significantly reduced in sporulation. The developmental defects of the five Δ*eps* mutants were restored by ectopic expression of the corresponding gene (Fig. 8).

**Figure 8.**
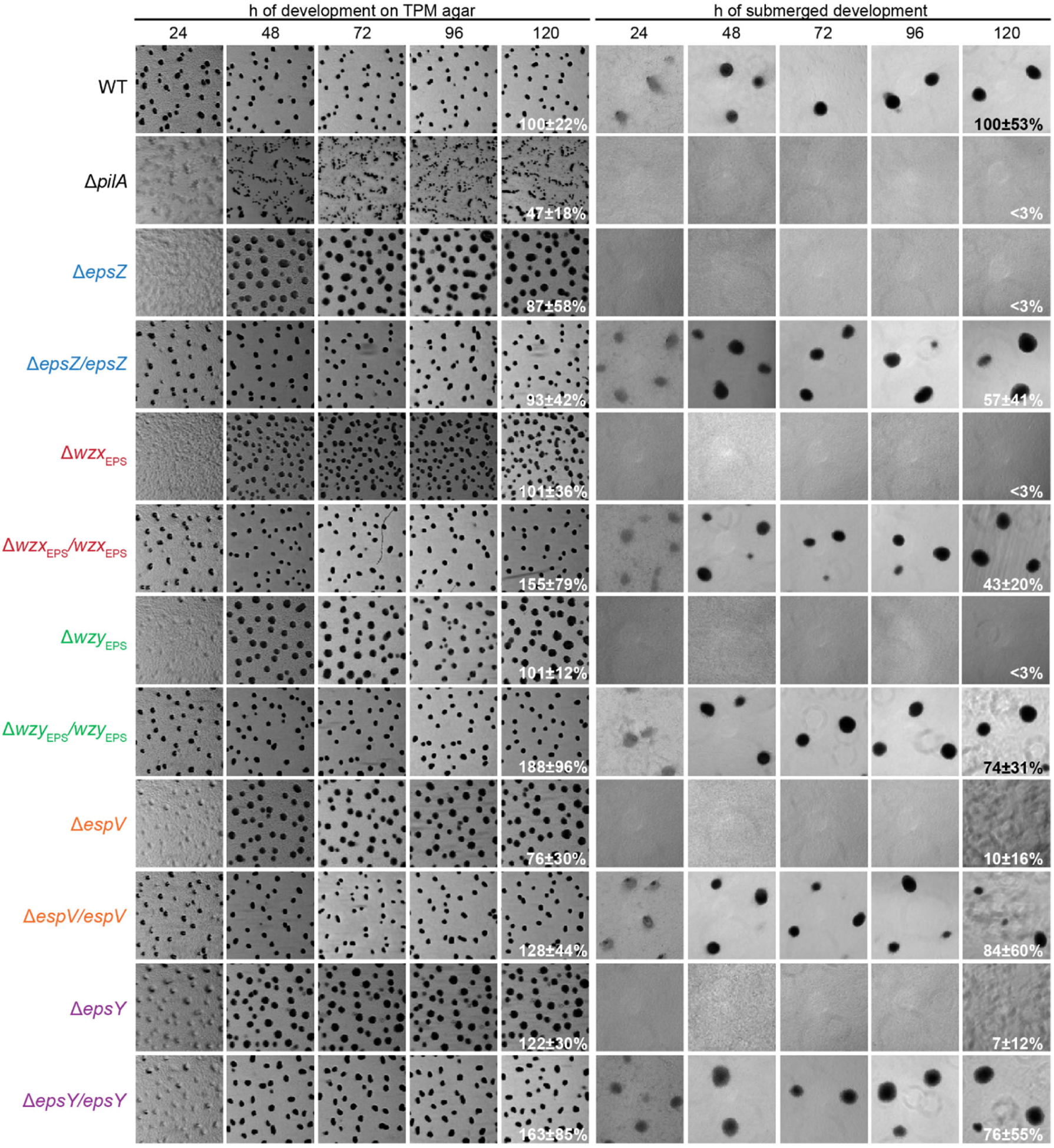
Development of Δ*eps* mutants. Cells on TPM agar and under submerged conditions were followed during development. Images were recorded at the indicated time points. Sporulation efficiency after heat and sonic treatment is indicated as the mean ± standard deviation from at three biological replicates relative to WT. Scale bars: 1mm (left), 200 µm (right).

## Discussion

Here, we focused on elucidating key steps of EPS biosynthesis, and determined functional consequences of the loss of the EPS biosynthetic machinery. The EPS structure is unknown; however, chemical analyses support that it contains at least *N*-acetyl-glucosamine (GlcNAc), Glc and Gal while data for other monosaccharides vary depending on the analysis (71, 72).

Using bioinformatics, we identified the genes for all the components of a Wzx/Wzy pathway in the *eps* locus. Our experimental results support a model in which these genes encode the EPS biosynthesis machinery (Fig. 9A) and that synthesis of the EPS repeat unit is initiated by the PHPT homolog EpsZ (MXAN_7415). We demonstrate in heterologous expression experiments that EpsZ is functionally similar to the Gal-1-P transferase WbaP_Se_, suggesting that Gal is the first sugar of the EPS repeat unit. The *eps* locus encodes five GTs and inactivation of each of these five genes (9, 34, 42) causes a loss of EPS synthesis or T4P-dependent motility (Fig. 2A). Therefore, we suggest that these five GTs add monosaccharides to build the repeat unit, which is then translocated across the IM by the Wzx_EPS_ flippase (MXAN_7416). The repeat units are polymerized by the Wzy_EPS_ polymerase (MXAN_7442) with the help of the PCP protein EpsV (MXAN_7421) to make the EPS polysaccharide. In the last step, the EPS polymer is transported to the surface through the OPX protein EpsY (MXAN_7417). EpsC (MXAN_7449) is a serine O-acetyltransferase homolog, which is important but not essential for EPS synthesis (9). As previously suggested for a paralog encoded by *exoN* (41), which is important for spore coat polysaccharide synthesis, MXAN_7449 could be involved in O-acetylation of precursors for EPS synthesis. Finally, the predicted glycosyl hydrolase EpsB (MXAN_7450) is also important but not essential for EPS synthesis (9) and its biochemical function remains to be characterized. Overall, our genetic and functional analyses support that the EPS biosynthesis machinery is exclusively dedicated to EPS biosynthesis and not involved in LPS O-antigen or spore coat polysaccharide biosynthesis.

**Figure 9.**
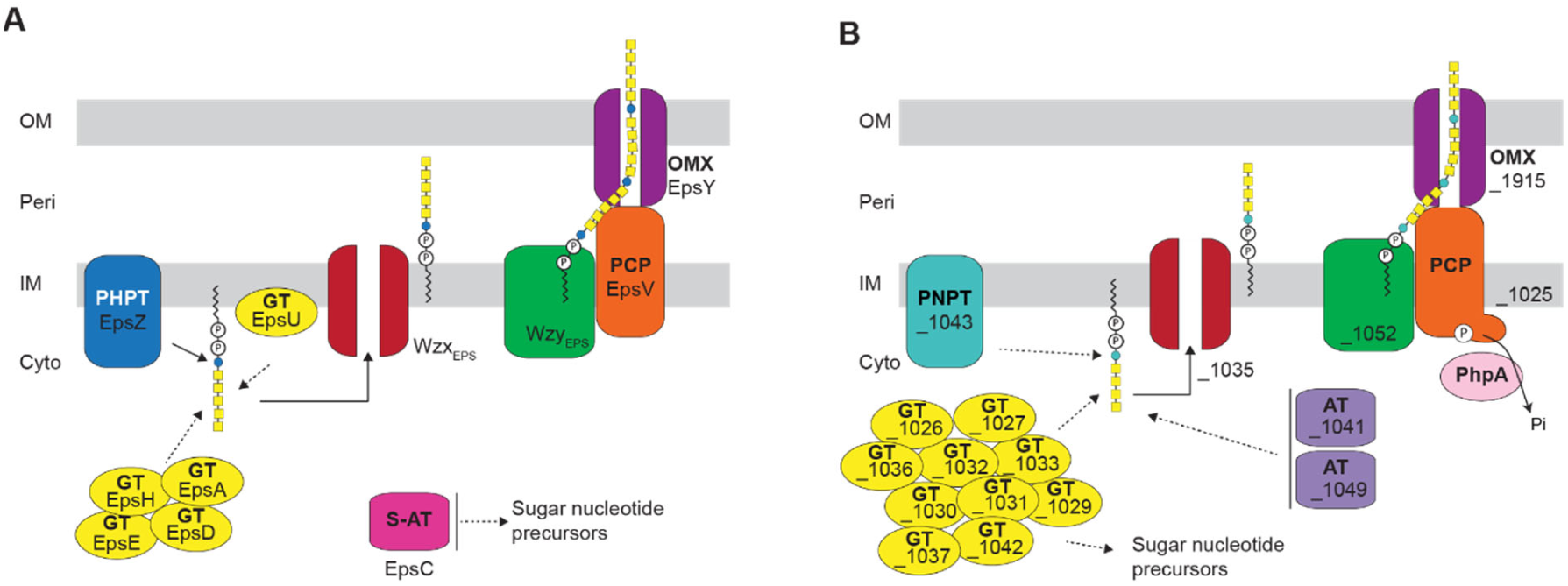
Wzx/Wzy-dependent pathways for EPS biosynthesis (A) and synthesis of an unknown polysaccharide (B). See text in Discussion for details.

We also identified two additional loci, which together encode a complete Wzx/Wzy-dependent pathway (Fig. 9B). Our genetic analysis suggests the proteins of this pathway are not involved in EPS biosynthesis, spore coat polysaccharide and LPS-O-antigen synthesis (Fig. 3D; unpublished data), indicating a novel function. A future goal will be to identify the polysaccharide synthesized by these proteins as well as its function. Islam et al. (73) have suggested these gene clusters encode a biosynthetic machinery for synthesis of a biosurfactant but further studies are required for its complete characterization.

Genetic analyses of the five core components of the EPS biosynthesis machinery showed that lack of any of these proteins caused a defect not only in EPS synthesis but also in T4P-dependent motility, cell-cell cohesion, and a conditional defect in fruiting body formation. Superficially, these defects are similar to those reported for *dif* mutants with an EPS^-^ phenotype, which are the best studied mutants with decreased EPS synthesis. However, more detailed comparisons reveal important differences. First, *dif* mutants with an EPS^-^ phenotype have a defect in T4P-dependent motility (17, 18); however, a *difA* mutant makes T4P at WT levels (21) or is hyper-piliated (15). Moreover, it was suggested that EPS stimulates T4P retractions in this mutant because addition of EPS caused reduced piliation (15, 19). Because the *dif* mutants with an EPS^-^ phenotype make T4P but have reduced T4P-dependent motility, this supports that EPS *per se* might stimulate T4P-dependent motility. By contrast, we observed that the five Δ*eps* mutants analyzed here are hypo-pilated. Also, further deletion of the gene for the PilT retraction ATPase resulted in an increased level of surface piliation suggesting that T4P in the five Δ*eps pilT*^*+*^ mutants can still be retracted. Consistently, T4P-dependent motility was not completely abolished in the five Δ*eps* mutants. These observation suggest that that EPS, or alternatively components of the EPS biosynthetic machinery, is important for T4P formation. Altogether, these comparisons support that the *dif* EPS^-^ mutations, which are regulatory mutants, and the Δ*eps* mutations described here, which are biosynthetic mutants, both interfere with T4P-dependent motility but the underlying mechanisms are different. Second, *dif* mutants with an EPS^-^ phenotype neither develop to form spore-filled fruiting bodies on TPM or CF agar nor under submerged conditions (17, 18, 43). Of note, development of such mutants on TPM agar was rescued by addition of EPS (21, 74). By contrast, the five Δ*eps* mutants described here develop with only a slight delay on TPM agar but not under submerged conditions. We speculate that this developmental defect is caused by lack of cell-cell cohesion and agglutination in the five Δ*eps* mutants. Whether these phenotypic differences are caused by the differences in T4P levels and functionality in the two types of mutants remains to be investigated.

Previously, it was reported that the T4P machinery functions upstream of the Dif pathway to stimulate EPS synthesis (75-78). How the T4P machinery interfaces with the Dif system is unknown. Similarly, it is unknown how the Dif system stimulates EPS biosynthesis. Here, we show that mutations in the Wzx/Wzy-dependent pathway for EPS synthesis cause a defect in T4P extension. It will be an important future goal to disentangle how *dif* and *eps* mutants at the molecular level affect T4P formation and function as well as how the T4P machinery affects EPS synthesis.

## Acknowledgement

The authors thank A. Treuner-Lange for construction of pMAT150. This work was supported by Deutsche Forschungsgemeinschaft (DFG, German Research Council) within the framework of the SFB987 “Microbial Diversity in Environmental Signal Response” as well as by the Max Planck Society.

## Data Availability

The data that support the findings of this study are available from the corresponding author upon request.

## Materials and Methods

### Strains and cell growth

All *M. xanthus* strains are derivatives of the wild type DK1622 (79). Strains, plasmids and oligonucleotides used in this work are listed in Table 1, Table 2, and Table S3, respectively. *M. xanthus* was grown at 32°C in 1% CTT (1% (w/v) Bacto Casitone, 10 mM Tris-HCl pH 8.0, 1 mM K_2_HPO_4_/KH_2_PO_4_ pH 7.6 and 8 mM MgSO_4_) liquid medium or on 1.5% agar supplemented with 1% CTT and kanamycin (50 µg ml^-1^) or oxytetracycline (10 µg ml^-1^), as appropriate (80). In-frame deletions were generated as described (81), and plasmids for complementation experiments were integrated in a single copy by site specific recombination into the Mx8 *attB* site. In-frame deletions and plasmid integrations were verified by PCR. Plasmids were propagated in *E. coli* Mach1 and DH5α.

**Table 1.**
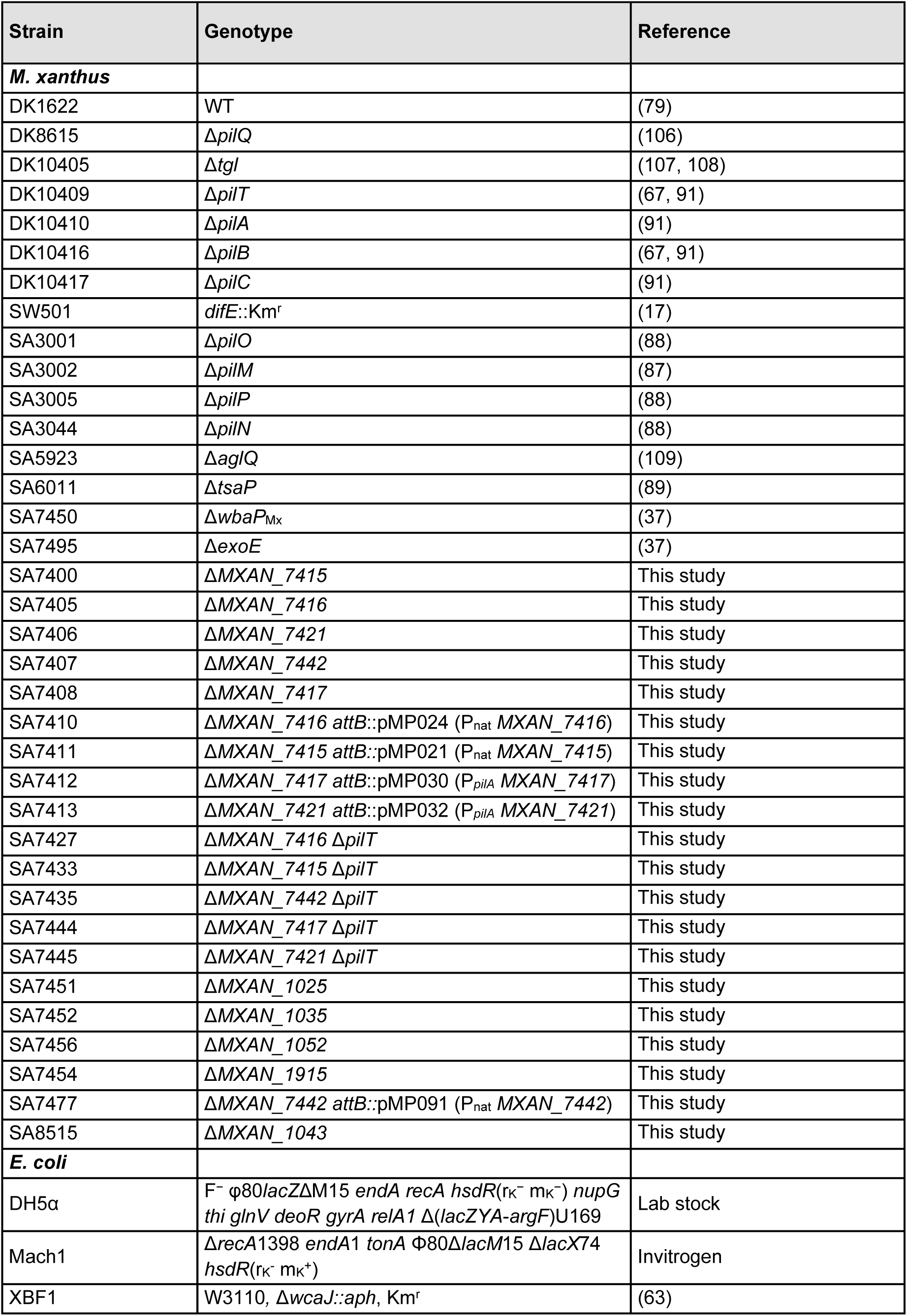
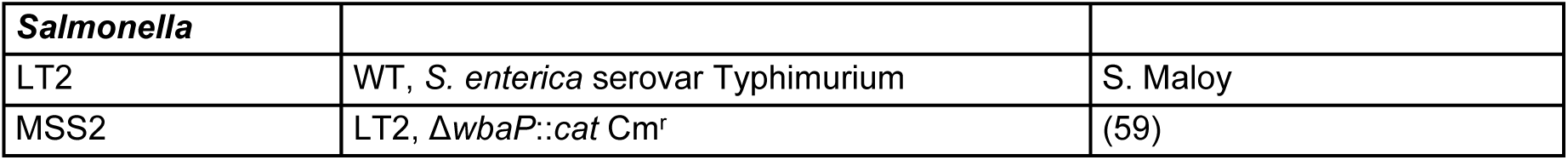
Strains used in this work

**Table 2.**
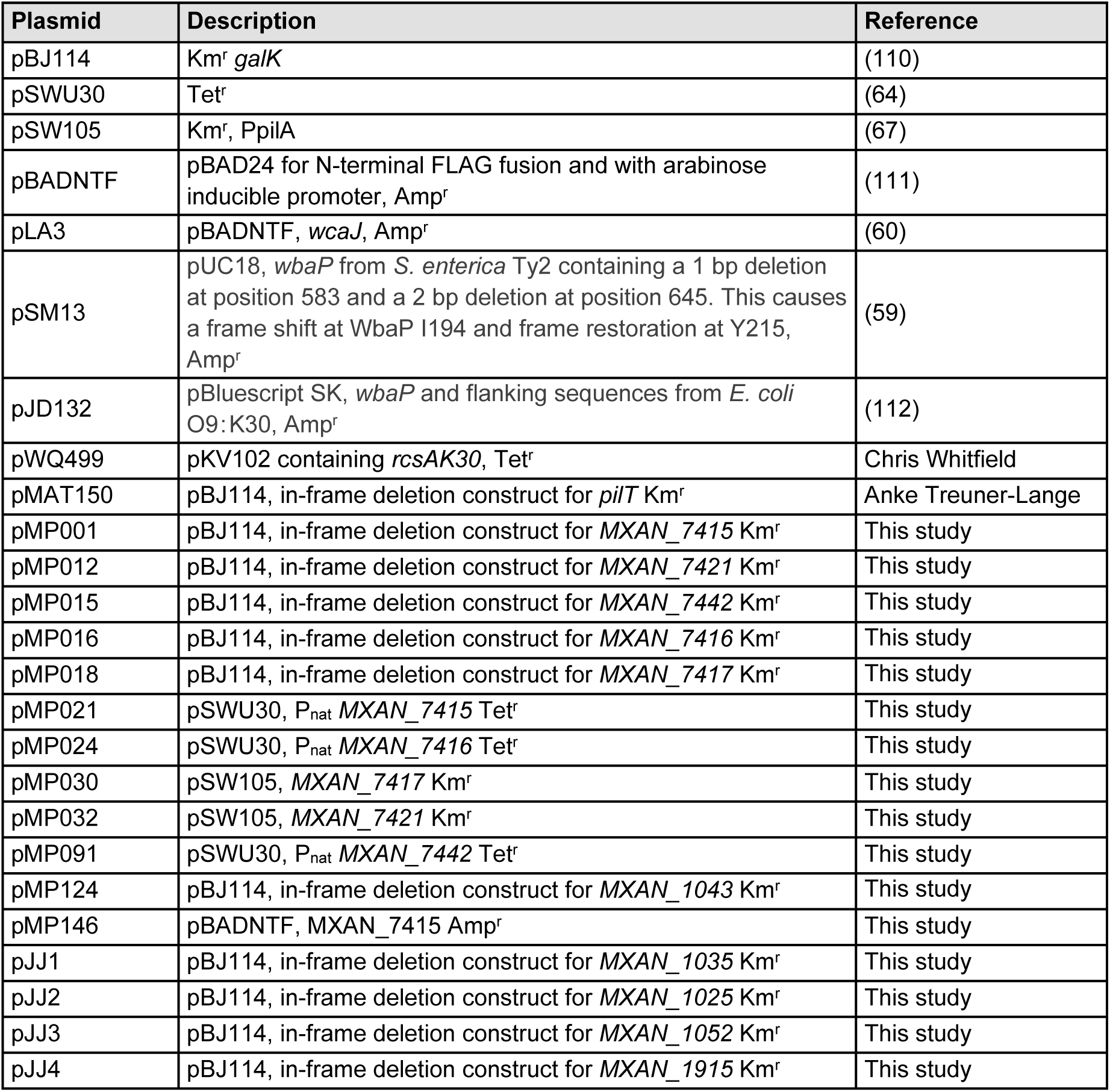
Plasmids used in this work

*E. coli* and *S. enterica* serovar Typhimurium strains were grown at 37°C in Luria-Bertani (LB) medium (10 mg tryptone ml^-1^, 5 mg yeast extract ml^-1^; 5 mg NaCl ml^-1^) supplemented, when required, with ampicillin, tetracycline, kanamycin or chloramphenicol at final concentrations of 100, 20, 40 and 30 µg ml^-1^, respectively. Plasmids for heterologous complementation were introduced into MSS2 and XBF1 strains (Table 1) by electroporation (82).

### Detection of EPS accumulation

Exponentially growing cells were harvested, (3 min, 6000 *g* at RT), and resuspended in 1% CTT to a calculated density of 7 × 10^9^ cells ml^-1^. 20 µl aliquots of the cell suspensions were placed on 0.5% agar plates supplemented with 0.5% CTT and 10 or 20 μg ml^-1^ of Trypan blue or Congo red, respectively. Plates were incubated at 32°C and documented at 24 h.

### Glycerol-induced sporulation assay

Sporulation in response to 0.5 M glycerol was performed as described (83) with a slightly modified protocol. Briefly, cells were cultivated in 10 ml of CTT medium, at a cell density of 3 × 10^8^ cells ml^-1^, glycerol was added to a final concentration of 0.5 M. At 0, 4 and 24 h after glycerol addition, cell morphology was observed by placing 5 µl of cells on a 1.5% agar TPM pad on a slide. Cells were immediately covered with a coverslip and imaged with DMi6000B microscope and a Hamamatsu Flash 4.0 Camera (Leica). To determine the resistance to heat and sonication of spores formed, cells from 5 ml of the culture after 24 h incubation were harvested (10 min, 4150 *g*, RT), resuspended in 1 ml sterile water, incubated at 50°C for 2 h, and then sonicated with 30 pulses, pulse 50%, amplitude 75% with a UP200St sonifier and microtip (Hielscher). Sporulation levels were determined as the number of sonication- and heat-resistant spores relative to WT using a Helber bacterial counting chamber (Hawksley, UK).5 μl of the treated samples were placed on a 1.5 % agar TPM pad on a slide, covered with a coverslip and imaged.

### LPS extraction and detection

LPS was extracted from *M. xanthus* and visualized by Emerald staining as described (37). LPS from *S. enterica* and *E. coli* was extracted and visualized by silver staining as described (37, 84). For *S. enterica*, O-antigen was detected by immunoblot using rabbit *Salmonella* O antiserum group B (Difco, Beckton Dickinson ref. number 229481) (1:500) and the secondary antibody IRDye 800CW goat α-rabbit immunoglobulin G (1:10000) (LI-COR) (37).

### Cell length determination

5 µl aliquots of exponentially growing cell suspensions were spotted on glass placed on a metal frame, covered with 1.5% agar supplemented with TPM and imaged using a DMi8 Inverted microscope and DFC9000 GT camera (Leica) (85). Cell length was determined and visualized as described (37). Statistical analyses were performed using SigmaPlot v14. All data sets were tested for a normal distribution using a Shapiro-Wilk test and for all data sets without a normal distribution, the Mann-Whitney test was applied to test for significant differences.

### Motility assays

Exponentially growing cultures of *M. xanthus* were harvested (6000 *g*, room temperature (RT)) and resuspended in 1% CTT to a calculated density of 7 × 10^9^ cells ml^-1^. 5 µl aliquots of cell suspensions were spotted on 0.5% and 1.5% agar supplemented with 0.5% CTT. The plates were incubated at 32°C for 24 h and cells were visualized using a M205FA Stereomicroscope (Leica) and imaged using a Hamamatsu ORCA-flash V2 Digital CMOS camera (Hamamatsu Photonics). Pictures were analyzed using Metamorph® v 7.5 (Molecular Devices).

### Detection of colanic acid biosynthesis

*E. coli* Δ*wcaJ* strains were grown on LB plates with antibiotics and with or without 0.2 % (w/v) arabinose at 37°C overnight. Incubation was extended to 24-48 h at RT to visualize the mucoid phenotype (Furlong et al 2015).

### Immunoblot analysis

Immunoblots were carried out as described (86). For *M. xanthus* immunoblots, rabbit polyclonal α-PilA (dilution: 1:2000), α-PilB (dilution: 1:2000) (67), α-PilC (dilution: 1:2000) (87), α-PilM (dilution: 1:3000) (87), α-PilN (dilution: 1:2000) (88), α-PilO (dilution: 1:2000) (88), α-PilP (dilution: 1:2000) (88), α-PilT (dilution: 1:3000) (67), α-Tgl (dilution: 1:2000) (88), α-TsaP (dilution: 1:2000) (89), α-PilQ (dilution: 1:5000) (87) were used together with a horseradish-conjugated goat anti-rabbit immunoglobulin G (Sigma) as secondary antibody. Blots were developed using Luminata crescendo Western HRP Substrate (Millipore) on a LAS-4000 imager (Fujifilm).

For *E. coli* and *S. enterica* strains, FLAG-tagged membrane proteins were isolated and detected by immunoblot analysis, as previously described, using α-FLAG M2 monoclonal antibody (Sigma) (1:10000) and a secondary antibody, IRDye 800CW Goat α-Mouse IgG (H+L), 0.5 mg (LI-COR) (1:10000) (37).

### T4P shear off assay

T4P were sheared from cells that had been grown for three days on 1.5% agar plates supplemented with 1% CTT at 32°C as described, except that precipitation of sheared T4P was done using TCA as described (90), and analyzed by immunoblotting with α-PilA antibodies as described previously (64). Blots were developed as indicated.

### Cell agglutination assay

Cell agglutination was performed as described previously (91) with a slightly modified protocol. Briefly, 1 ml of exponentially growing cells in 1% CTT was transferred to a cuvette and cell density was measured at the indicated time points.

### Development

Exponentially growing *M. xanthus* cultures were harvested (3 min, 6000 *g* at RT), and resuspended in MC7 buffer (10 mM MOPS pH 7.0, 1 mM CaCl2) to a calculated density of 7 × 10^9^ cells ml^-1^. 10 μl aliquots of cells were placed on TPM agar (10 mM Tris-HCl pH 7.6, 1 mM K_2_HPO_4_/KH_2_PO_4_ pH 7.6, 8 mM MgSO_4_), and 50 μl aliquots were mixed with 350 μl of MC7 buffer and placed in a 24-well polystyrene plate (Falcon) for development in submerged culture. Cells were visualized at the indicated time points using a M205FA Stereomicroscope (Leica) and imaged using a Hamamatsu ORCA-flash V2 Digital CMOS camera (Hamamatsu Photonics), and a DMi8 Inverted microscope and DFC9000 GT camera (Leica). Images were analyzed as previously described. After 120 h, cells were collected and incubated at 50°C for 2 h, and then sonicated as described above. Sporulation levels were determined as the number of sonication- and heat-resistant spores relative to WT.

### Bioinformatics

The KEGG SSDB (Sequence Similarity Database) (92) database was used to identify homologs of PHPT (PF02397-Bacterial Sugar Transferase), PNPT (PF00953-Glycosyl transferase family 4) (93), Wzx (PF01943- Polysacc_synt and PF13440- Polysacc_synt_3), Wzy_C (PF04932- Wzy_C), PCP (PF02706- Wzz) and OPX (PF02563- Poly_export) as in (41, 94, 95). For the ABC-transporter dependent pathway we used (PF01061-ABC2_membrane) for the permease and, (PF00005- ABC_tran) and (PF14524- Wzt_C) for the ATPase as in (37) together with an analysis of the genetic neighborhood to search for glycan related proteins. BlastP was used to identify homologs of the synthase dependent pathway using previously identified components (33). KEGG SSDB was also used to identify EPS homolog proteins in other Myxococcales using a reciprocal best BlastP hit method. UniProt (96), KEGG (92) and the Carbohydrate Active Enzymes (CAZy) (http://www.cazy.org/) (97) databases were used to assign functions to proteins (Fig. 1B, 2A, 3A-B; Table S1 and S2).SMART (smart.embl-heidelberg.de) (98) and Pfam v31.0 and v32.0 (pfam.xfam.org) (99) were used to identify protein domains. Membrane topology was assessed by TMHMM v2.0 (100) and two-dimensional topology was graphically shown using TOPO2 (101). Clustal Omega (102) was used to align protein sequences. The phylogenetic tree was prepared as in (41) in MEGA7 (103) using the Neighbor-Joining method (104). Bootstrap values (500 replicates) are shown next to the branches (105).

