## Supplementary Information for "Characterization of the exopolysaccharide biosynthesis pathway in *Myxococcus xanthus*"

**This file contains:**

Supplementary Figure 1

Supplementary Experimental Procedures

Table S1-S3

Supplementary References

### Supplemental Figures & Legends

```
WbaPSe  ---------- ---------- ---MDNIDNK YNPQLCKIFL AISDLIFFNL ALWFSLGCVY FIFDQVQRFI PQDQLDTRVI  57
EpsZ    MDTMSGATAS AAVVGAPGSA NAQGQVIEEV QGPPKLAPGS AAKLNLTVDL VLLSSVLVGS AWLSGQLSAE SGWKVSGLVL  80

WbaPSe  THFILSVVCV GWFWIRLRHY TYRKPFWYEL KEIFRTIVIF AIFDLALIAF TKWQFSRYVW VFCWTFALIL VPFFRALTKH  137
EpsZ    AAWVVWIVTG TALCLYDSRF AERSKLDHVA LVSVTTLAVV TVLTLGSVAV PTVVTSPVVG PLLFIFWPVT LLLRLFVFRP  160

WbaPSe  LLNKLGIWKK KTIILGSGQN ARGAYSALQS EEMMGFDVIA FFDTDASDAE INMLP--VIK DTEIIWDLNR --TGDVHYIL  213
EpsZ    VASQE---RP MDAVLIVGTG AMGRYTGEDL ANRGRRQILG YVRFHDDNGS VGELPGPVMG SVDDLEHILR NTAVDEVYIA  237

WbaPSe  AYEYTELEKT HFWLRELSKH HCRSVTVVPS FRGLPLYNTD MSFIFSHEVM LLRIQNNLAK RSSRFLKRTF DIVCSIMILI  293
EpsZ    GNTLKQGESM QAAIKLAERF GVPFALPAHS FRLDRARPVE RRAVADGFLH FAAVS---PK PHQMAMKRLF DICVSAAALW  314

WbaPSe  IASPLMIYLW YKVTRDG-GP AIYGHQRVGR HGKLFPCYKF RSMVMNSQEV LKELLANDPI ARAEWEKDFK LKNDPRITAV  372
EpsZ    ALLPLLGMVA LAVKFTSKGP IFFKQLRVGQ NGKPFYMLKF RSMVVNAEEL KEKLAALN-- --EQTGPVFK MKHDPRITGI  390

WbaPSe  GRFIRKTSLD ELPQLFNVLK GDMSLVGPRP IVSDELERYC DDVDYYLMAK PGMTGLWQVS GRNDVDYDTR VYFDSWYVKN  452
EpsZ    GRFIRKFSID ELPQFINVLR GEMSIVGPRP PVPTEVAKYE TWQRRRLSVR PGLTCIWQVS GRNQISFEEW MYLDMQYIDH  470

WbaPSe  WTLWNDIAIL FKTAKVVLRR DGAY  476
EpsZ    WSLTSDLRLL LQTVPVVLTG RGAS  494
```

**Figure S1.** Sequence alignment of EpsZ (MXAN_715) and $WbaP_{Se}$ (Accession number: NP_461027.1) showing the Pro and Asp (orange) residues in the motif $DX_{12}P$ and the conserved amino acids essential for catalytic activity (red).

**Supplementary Experimental Procedures**

Plasmid construction. All oligonucleotides used are listed in Table S3. All constructed plasmids were verified by DNA sequencing.

**pMP001** (for generation of in-frame deletion of *MXAN_7415*): up- and downstream fragments were amplified from genomic DNA of DK1622 using the primer pairs 7415_A/7415_B and 7415_C/7415_D respectively, as described in (1). Subsequently, the AB and CD fragments were used as templates to perform an overlapping PCR with the primer pair 7415_A/7415_D to generate the AD fragment. The AD fragment was digested with KpnI/XbaI and cloned in pBJ114.

**pMP012** (for generation of in-frame deletion of *MXAN_7421*): up- and downstream fragments were amplified from genomic DNA of DK1622 using the primer pairs 7421_A/7421_B and 7421_C/7421_D respectively. Subsequently, the AB and CD fragments were used as templates to perform an overlapping PCR with the primer pair 7421_A/7421_D to generate the AD fragment. The AD fragment was digested with KpnI/XbaI and cloned in pBJ114.

**pMP015** (for generation of in-frame deletion of *MXAN_7442*): up- and downstream fragments were amplified from genomic DNA of DK1622 using the primer pairs 7442_A/7442_B and 7442_C/7442_D respectively. Subsequently, the AB and CD fragments were used as templates to perform an overlapping PCR with the primer pair 7442_A/7442_D to generate the AD fragment. The AD fragment was digested with KpnI/XbaI and cloned in pBJ114.

**pMP016** (for generation of in-frame deletion of *MXAN_7416*): up- and downstream fragments were amplified from genomic DNA of DK1622 using the primer pairs 7416_A/7416_B and 7416_C/7416_D respectively. Subsequently, the AB and CD fragments were used as templates to perform an overlapping PCR with the primer pair 7416_A/7416_D to generate the AD fragment. The AD fragment was digested with KpnI/XbaI and cloned in pBJ114.

**pMP018** (for generation of in-frame deletion of *MXAN_7417*): up- and downstream fragments were amplified from genomic DNA of DK1622 using the primer pairs 7417_A/7417_B and 7417_C/7417_D respectively. Subsequently, the AB and CD fragments were used as templates to perform an overlapping PCR with the primer pair 7417_A/7417_D to generate the AD fragment. The AD fragment was digested with EcoRI/XbaI and cloned in pBJ114.

**pMP021** (expression of $P_{nat}$ *MXAN_7415* from the *attB* site): $P_{\mathrm{nat}}$ *MXAN_7415* was amplified with the primer combination 7415 Pnat900 forw2/7415 Pnat rev and genomic DNA from *M. xanthus*

DK1622 as a template. The fragment was digested with KpnI and XbaI, cloned into pSWU30 and sequenced.

**pMP024** (expression of $P_{nat}$ *MXAN_7416* from the *attB* site): $P_{nat}$ *MXAN_7416* was amplified with the primer combination 7416_Pnat forw2/7416_Pnat rev and genomic DNA from *M. xanthus* DK1622 as a template. The fragment was digested with HindIII and XbaI, cloned into pSWU30 and sequenced.

**pMP030** (expression of $P_{pilA}$ *MXAN_7417* from the *attB* site): *MXAN_7417* was amplified with the primer combination 7417_PpilA forw/7417_PpilA rev and genomic DNA from *M. xanthus* DK1622 as a template. The fragment was digested with XbaI and HindIII, cloned into pSW105 and sequenced.

**pMP032** (expression of $P_{pilA}$ *MXAN_7421* from the *attB* site): *MXAN_7421* was amplified with the primer combination 7421_PpilA forw/7421_PpilA rev and genomic DNA from *M. xanthus* DK1622 as a template. The fragment was digested with XbaI and HindIII, cloned into pSW105 and sequenced.

**pMP091** (expression of $P_{nat}$ *MXAN_7442* from the *attB* site): $P_{nat}$ *MXAN_7442* was amplified with the primer combination 7442_Pnat 600 up /7442_Pnat rev and genomic DNA from *M. xanthus* DK1622 as a template. The fragment was digested with HindIII and XbaI, cloned into pSWU30 and sequenced.

**pMP124** (for generation of in-frame deletion of *MXAN_1043*): up- and downstream fragments were amplified from genomic DNA of DK1622 using the primer pairs 1043_A/1043_B and 1043_C/1043_D respectively. Subsequently, the AB and CD fragments were used as templates to perform an overlapping PCR with the primer pair 1043_A/1043_D to generate the AD fragment. The AD fragment was digested with EcoRI/HindIII and cloned in pBJ114.

**pMP146** (expression of *MXAN_7415* under the control of an arabinose promoter): *MXAN_7415* was amplified with the primer combination 7415 fw +1/ 7415 rev new (pilA) and genomic DNA from *M. xanthus* DK1622 as a template. The fragment was digested with XbaI and HindIII, cloned into pBADNTF and sequenced.

**pJJ1** (for generation of in-frame deletion of *MXAN_1035*): up- and downstream fragments were amplified from genomic DNA of DK1622 using the primer pairs 1035_A/1035_B and 1035_C/1035_D respectively. Subsequently, the AB and CD fragments were used as templates to perform an overlapping PCR with the primer pair 1035_A/1035_D to generate the AD fragment. The AD fragment was digested with KpnI/XbaI and cloned in pBJ114.

**pJJ2** (for generation of in-frame deletion of *MXAN_1025*): up- and downstream fragments were amplified from genomic DNA of DK1622 using the primer pairs 1025_A/1025_B and 1025_C/1025_D respectively. Subsequently, the AB and CD fragments were used as templates to perform an overlapping PCR with the primer pair 1025_A/1025_D to generate the AD fragment. The AD fragment was digested with KpnI/XbaI and cloned in pBJ114.

**pJJ3** (for generation of in-frame deletion of *MXAN_1052*): up- and downstream fragments were amplified from genomic DNA of DK1622 using the primer pairs 1025_A/1025_B and 1052_C/1052_D respectively. Subsequently, the AB and CD fragments were used as templates to perform an overlapping PCR with the primer pair 1052_A/1052_D to generate the AD fragment. The AD fragment was digested with KpnI/BamHI and cloned in pBJ114.

**pJJ4** (for generation of in-frame deletion of *MXAN_1915*): up- and downstream fragments were amplified from genomic DNA of DK1622 using the primer pairs 1915_A/1915_B and 1915_C/1915_D respectively. Subsequently, the AB and CD fragments were used as templates to perform an overlapping PCR with the primer pair 1915_A/1915_D to generate the AD fragment. The AD fragment was digested with KpnI/XbaI and cloned in pBJ114.

**pMAT150** (for generation of in-frame deletion of *pilT*): up- and downstream fragments were amplified from genomic DNA of DK10409 (Δ*pilT*) using the primer pairs pilT-A EcoRI/ pilT-D HindIII. The AD fragment was digested with EcoRI/HindIII and cloned in pBJ114.

**Table S1.** Analysis of the *eps* locus

| Locus tag MXAN | Gene name | (Putative) function of encoded protein | Reference[1] |
|---|---|---|---|
| *7415* | *epsZ* | Polyprenyl glycosylphosphotransferase<br>**New annotation: polyisoprenyl-phosphate hexose-1-phosphate transferase** | (2) |
| *7416* | ***wzx***$_{EPS}$ | Wzx flippase | (3) |
| *7417* | *epsY* | Polysaccharide biosynthesis/export protein<br>**New annotation: OPX protein** | Uniprot, KEGG |
| *7418* | *epsX* | Hypothetical protein | |
| *7420* | *epsW* | Response regulator | (4) |
| *7421* | *epsV* | Chain length determinant protein<br>**New annotation: Wzz protein** | Uniprot, KEGG |
| *7422* | *epsU* | Glycosyltransferase | |
| *7423* | | Hypothetical protein | |
| *7424* | | Hypothetical protein | |
| *7425* | | Hypothetical protein | |
| *7426* | | Hypothetical protein | |
| *7430* | | Transposase orfB, IS5 family | Uniprot, KEGG |
| *7431* | *epsP* | Transposase orfA, IS5 family | |
| *7433* | *epsO* | von Willebrand factor type A domain protein, Ca-activated chloride channel homolog | |
| *7435* | *epsN* | Hydrolase | |
| *7436* | *epsM* | Outer membrane efflux protein, cobalt-zinc-cadmium efflux system | |
| *7437* | *epsL* | Heavy metal efflux pump, CzcA family, cobalt-zinc-cadmium resistance protein | Uniprot, KEGG |
| *7438* | *epsK* | putative cobalt-zinc-cadmium resistance protein | Uniprot, KEGG |
| *7439* | *epsJ* | Sensor histidine kinase | |
| *7440* | *epsI*<br>*nla24* | Sigma-54 dependent DNA-binding response regulator | (5) |
| *7441* | *epsH* | Glycosyltransferase | |
| *7442* | *sgnF*,<br>***wzy***$_{EPS}$ | Putative membrane protein, Wzy_C domain<br>**New annotation: Wzy polymerase** | (6) |
| *7443* | *epsG* | Magnesium transporter | |
| *7444* | *epsF* | Response regulator/sensory box histidine kinase | |
| *7445* | *epsE* | Glycosyltransferase | |
| *7447* | | Hypothetical protein | Uniprot, KEGG |
| *7448* | *epsD* | Glycosyltransferase | |
| *7449* | *epsC* | Serine O-acetyltransferase | |
| *7450* | *epsB* | Glycosyl hydrolase | Uniprot, KEGG |
| *7451* | *epsA* | Glycosyltransferase | Uniprot, KEGG |

[1]Based on (7) unless indicated otherwise.

**Table S2.** Analysis of the *MXAN_1025-MXAN_1052* and *MXAN_1915* loci

| Locus tag MXAN | Gene name | (Putative) function of encoded protein | Reference[1] |
|---|---|---|---|
| *1025* | | Bacterial tyrosine kinase, Capsular exopolysaccharide family protein<br>**New annotation: Wzc** | (8) |
| *1026* | | Glycosyltransferase | |
| *1027* | | Glycosyltransferase | |
| *1028* | | Putative membrane protein | |
| *1029* | | Glycosyltransferase | |
| *1030* | | Glycosyltransferase | |
| *1031* | | Glycosyltransferase | |
| *1032* | | Glycosyltransferase | |
| *1033* | | Glyco_trans_4-like_N domain-containing protein | |
| *1034* | | Conserved domain protein | |
| *1035* | | Putative membrane protein, PST family<br>**New annotation: Wzx flippase** | |
| *1036* | | Glycosyltransferase | |
| *1037* | | Glycosyltransferase | |
| *1038* | | Hypothetical protein | |
| *1039* | *glkA* | Glucokinase | |
| *1040* | | Sulfatase family protein | |
| *1041* | | Acyltransferase family protein | |
| *1042* | | Glycosyltransferase | |
| *1043* | | Glycosyltransferase<br>**New annotation: polyisoprenyl-phosphate *N*-acetylhexosamine-1-phosphate transferase** | |
| *1045* | | Hypothetical protein | |
| *1046* | | FG-GAP repeat/HVR domain protein | |
| *1047* | | Hypothetical protein | |
| *1048* | | UDP-glucose 6-dehydrogenase | |
| *1049* | | Acyltransferase family protein | |
| *1050* | | Hypothetical protein | |
| *1051* | | Hypothetical protein | |
| *1052* | | O-antigen polymerase family protein<br>**New annotation: Wzy polymerase** | |
| *1914* | *suhB* | Inositol-1-monophosphatase | |
| *1915* | | Polysaccharide biosynthesis/export protein<br>**New annotation: OPX protein** | |
| *1916* | | Hypothetical protein | |
| *1917* | | Hypothetical protein | |

[1]Based on Uniprot and KEGG, unless indicated otherwise.

**Table S3.** Oligonucleotides used in this work[1]

| Primer name | Sequence 5‘-3‘ | Brief descrption |
|---|---|---|
| 7415_A | ATCGGGTACCGTGGTGCTCGCCGTCAGTGG | For Δ*MXAN_7415* |
| 7415_B | CACCGGCACCGGGGCCAGCTTGGGCGG | For Δ*MXAN_7415* |
| 7415_C | CTGGCCCCGGTGCCGGTGGTGCTCACG | For Δ*MXAN_7415* |
| 7415_D | ATCGTCTAGACCCCCGCCCCACACCAGCTT | For Δ*MXAN_7415* |
| 7415_E | ACCTCCTGGCCGCCCATGAG | For Δ*MXAN_7415* |
| 7415_F | CTTCACCGCCTCGGACGCCA | For Δ*MXAN_7415* |
| 7415_G | CATCTTCTGGCCGGTGACGC | For Δ*MXAN_7415* |
| 7415_H | GCATGTAGAAGGGCTTGCCG | For Δ*MXAN_7415* |
| 7415 Pnat900 forw2 | ATCGGGTACCTGAGCCTTCCTCGACGTGGAGCG | For complementation fw |
| 7415 Pnat rev | ATCGTCTAGACTAGCTGGCGCCGCGGCCCG | For complementation rev |
| 7415 fw +1 | ATCGTCTAGAGGTGGACACGATGAGCGGCGC | For protein expression under an arabinose inducible promoter fw. |
| 7415 rev new | ATCGAAGCTTCTAGCTGGCGCCGCGGCCCG | For protein expression under an arabinose inducible promoter rev. |
| 7416_A | ATCGGGTACCGCAGCTTCGCGTGGGGCAGA | For Δ*MXAN_7416* |
| 7416_B | TTCGGGCGCTTGGAGCCCGTTGCGCAC | For Δ*MXAN_7416* |
| 7416_C | GGGCTCCAAGCGCCCGAAGCCGCGCCC | For Δ*MXAN_7416* |
| 7416_D | ATCGTCTAGAGCGCCCCTCGGCGTGGATGA | For Δ*MXAN_7416* |
| 7416_E | TGAAGTTCACCTCCAAGGGC | For Δ*MXAN_7416* |
| 7416_F | TCCACCACCACCACGTCACC | For Δ*MXAN_7416* |
| 7416_G | CGAGGTGCGCCAGCTCGTCT | For Δ*MXAN_7416* |
| 7416_H | CCCATCAGCCCCACCCACAG | For Δ*MXAN_7416* |
| 7416_Pnat forw2 | ATCGAAGCTTTGACAAGCCTCCAGGCAACCCAA | For complementation fw |
| 7416_Pnat rev | ATCGTCTAGATCACGGGGTGGGCGCGGCTT | For complementation rev |
| 7417_A | ATCGGAATTCGATGATGCTCATCGTCCTGG | For Δ*MXAN_7417* |
| 7417_B | GTCACCGGGGCGGTGCGTGGACGGCAT | For Δ*MXAN_7417* |
| 7417_C | ACGCACCGCCCCGGTGACGTGGTGGTG | For Δ*MXAN_7417* |
| 7417_D | ATCGTCTAGAGCCGCTGATGGAGAAGCCGC | For Δ*MXAN_7417* |
| 7417_E | GCGCCGCTCGCTGGAGGGCA | For Δ*MXAN_7417* |
| 7417_F | CGCGGGCGGGCCCGTCCAGG | For Δ*MXAN_7417* |
| 7417_G | CGTCCCTGGCGCTCGTTCGC | For Δ*MXAN_7417* |
| 7417_H | CGCAGGCGGAAGGTGGGCGC | For Δ*MXAN_7417* |
| 7417_PpilA forw | ATCGTCTAGAGTGAGGAGAGTTCCACCGCT | For complementation fw |
| 7417_PpilA rev | ATCGAAGCTTTTATTCCACCACCACCACGT | For complementation rev |
| 7421_A | ATCGGGTACCCCCTGCCAGCCAAGGCGGCG | For Δ*MXAN_7421* |

| | | |
|---|---|---|
| 7421_B | CGCCAGCACGGGAGCCCCGGGCGCGGG | For Δ*MXAN_7421* |
| 7421_C | GGGGCTCCCGTGCTGGCGGAGCTGGAG | For Δ*MXAN_7421* |
| 7421_D | ATCGTCTAGAGCCAGGACGCCGTGGGGTTC | For Δ*MXAN_7421* |
| 7421_E | CGTGCGGCAAACTGGTATTC | For Δ*MXAN_7421* |
| 7421_F | AGGGCAATGGTCATCAGCCG | For Δ*MXAN_7421* |
| 7421_G | GAGCGCCCGGAGACGAACGC | For Δ*MXAN_7421* |
| 7421_H | CGCGAAGATGCCCATGCCGA | For Δ*MXAN_7421* |
| 7421_PpilA forw | ATCGTCTAGAGTGACGGTCCCCGCGCCCGG | For complementation fw |
| 7421_PpilA rev | ATCGAAGCTTTCAGCGCCGCTCCAGCTCCG | For complementation rev |
| 7442_A | ATCGGGTACCGGGCAGACCGCCATTGAGCG | For Δ*MXAN_7421* |
| 7442_B | GGAGGATGGGGCCAACGCGACCACGGG | For Δ*MXAN_7421* |
| 7442_C | GCGTTGGCCCCATCCTCCGCCGCGAAC | For Δ*MXAN_7421* |
| 7442_D | ATCGTCTAGACCAGAAGGTGGGTGGCACGG | For Δ*MXAN_7421* |
| 7442_E | CCACTCCTTCTCCCGCCGCC | For Δ*MXAN_7421* |
| 7442_F | ATCGTCAGCGTGGTGCAGGC | For Δ*MXAN_7421* |
| 7442_G | CGTGGGGGTGGTGTGGGTCA | For Δ*MXAN_7421* |
| 7442_H | CCTCGTTGGGGTAGGTGATG | For Δ*MXAN_7421* |
| 7442_Pnat 600 up | ATCGAAGCTTTGAGCCCTTGGGCCAGGGCAGAC | For complementation fw |
| 7442_Pnat rev | ATCGTCTAGATCAGCGCGGGTTCGCGGCGG | For complementation rev |
| 1025_A | ATAGGGTACCGTGACGGAGCGCAGCGCCTC | For Δ*MXAN_1025* |
| 1025_B | ATCCTTGGAGGCCGGGTCGAAACCGGT | For Δ*MXAN_1025* |
| 1025_C | GACCCGGCCTCCAAGGATGGGGTGGCG | For Δ*MXAN_1025* |
| 1025_D | ACTGTCTAGAAGTAGACGAGCCGCCCCACC | For Δ*MXAN_1025* |
| 1025_E | CTGCGCGGCCAGCTTCACCA | For Δ*MXAN_1025* |
| 1025_F | ACCCGGCGCAGGGCCTGTAG | For Δ*MXAN_1025* |
| 1025_G | GACGCGGTGGCGCTGGTCCA | For Δ*MXAN_1025* |
| 1025_H | ACGAAGAGGCCGGGCACCTC | For Δ*MXAN_1025* |
| 1035_A | ATCGGGTACCAGCCGGAGCGGTGCACCTGG | For Δ*MXAN_1035* |
| 1035_B | CGCCGGAGTGGCTTCGGGCGCTGGGGT | For Δ*MXAN_1035* |
| 1035_C | CCCGAAGCCACTCCGGCGAGTCCGGCG | For Δ*MXAN_1035* |
| 1035_D | ATCGTCTAGACTTCCAGGCCCGCACGCACC | For Δ*MXAN_1035* |
| 1035_E | GCTTCCAGCCGCTCATGCCG | For Δ*MXAN_1035* |
| 1035_F | TAATCACCGCCTCCGGGCAG | For Δ*MXAN_1035* |
| 1035_G | GCGGCCTCCCTGGGCGTGTT | For Δ*MXAN_1035* |
| 1035_H | AGCGCCTGCGCCACCACACC | For Δ*MXAN_1035* |
| 1043_A | ATCGGAATTCCATGCCGAAGGCGTGTCCTT | For Δ*MXAN_1043* |
| 1043_B | CAGGCGTCCGAAGAAGGCGACCAGAAG | For Δ*MXAN_1043* |
| 1043_C | GCCTTCTTCGGACGCCTGGTCGCGATG | For Δ*MXAN_1043* |
| 1043_D | ATCGAAGCTTCCCTCCAGCGCCTCGCCCCA | For Δ*MXAN_1043* |
| 1043_E | CGCGGAGATGGTGGCCGTCG | For Δ*MXAN_1043* |
| 1043_F | CGTCTCCGCCCGCGCCAGCA | For Δ*MXAN_1043* |
| 1043_G | TGGGGATGGCTGGACCAGGC | For Δ*MXAN_1043* |

| | | |
|---|---|---|
| 1043_H | TGTTGGCGAAGTTCAGCGCC | For Δ*MXAN_1043* |
| 1052_A | ATCGGGTACCTTGATGGAGCAGTCGCACAG | For Δ*MXAN_1052* |
| 1052_B | CTCGCCATCAGAGGCACTGCGGGAAGC | For Δ*MXAN_1052* |
| 1052_C | AGTGCCTCTGATGGCGAGGCTCGGGAC | For Δ*MXAN_1052* |
| 1052_D | ATCGGGATCCCTCGAACCCCTGTACGGCGC | For Δ*MXAN_1052* |
| 1052_E | GCGAAGGCACAGCTCCTTCT | For Δ*MXAN_1052* |
| 1052_F | GGATGAAGACGGGAGAGCGC | For Δ*MXAN_1052* |
| 1052_G | GCTCGTCGTGGTGTGTCCGC | For Δ*MXAN_1052* |
| 1052_H | CGAAGAACGCATCCGCGCCA | For Δ*MXAN_1052* |
| 1915_A | ATCGGGTACCGCTTCCTGCCGAAGACGGCG | For Δ*MXAN_1915* |
| 1915_B | CACGAAGACGGTGAGGGCGGCGCGGAA | For Δ*MXAN_1915* |
| 1915_C | GCCCTCACCGTCTTCGTGCCGGAGAGC | For Δ*MXAN_1915* |
| 1915_D | ATCGTCTAGACAGGGCACCGACAGCCTGCG | For Δ*MXAN_1915* |
| 1915_E | GACTACGGCAACCTGCGAGT | For Δ*MXAN_1915* |
| 1915_F | CCCCGACCACGCCATCCTCG | For Δ*MXAN_1915* |
| 1915_G | TGACGCTGCCCGCCTGCTTC | For Δ*MXAN_1915* |
| 1915_H | TCGCCGGGCTGGAGCATGAA | For Δ*MXAN_1915* |
| pilT-A EcoRI | GCGCGAATTCCGCGACTTCGAGACGGCGG | For Δ*pilT* |
| pilT-D HindIII | GCGCAAGCTTGAGCTTCTCGTTCTTCTCC | For Δ*pilT* |
| pilT_E | CTCCGCCAGGACCCGGACATC | For Δ*pilT* |
| pilT_F | TATCGAGGCACTGCACCA | For Δ*pilT* |
| pilT_G | CTTGAAGACGGCGCCGCTGA | For Δ*pilT* |
| pilT_H | CGCGCTGATTCACGAGGCAG | For Δ*pilT* |

[1] Underlined sequences indicate restriction sites.
